# Epigenetic Regulation of Inflammation by Dopamine in Primary Human Macrophages

**DOI:** 10.64898/2026.01.21.700899

**Authors:** Yash Agarwal, Margish Ramani, Samyuktha Manikandan, Kimberly Bonar, John Montilla, Peter J. Gaskill, Stephanie M. Matt

**Affiliations:** Drexel University College of Medicine, Department of Pharmacology and Physiology, Philadelphia, PA, USA; Drexel University College of Medicine, Department of Medicine, Philadelphia, PA, USA

**Author notes:** Correspondence: Stephanie M. Matt, Ph.D. Department of Pharmacology and Physiology Drexel University College of Medicine 245 N. 15th Street, Philadelphia, PA, 19102. Co-last authors (Peter J. Gaskill, Stephanie M. Matt).

**Keywords:** DNA methylation, Dopamine, Epigenetics, Inflammation, Macrophage

## Abstract

While dopamine is a monoamine neurotransmitter best known for its roles in reward, motivation, and motor function in the central nervous system, its actions extend beyond neurons and can influence non-neuronal cells via epigenetic mechanisms. An increasing body of literature corroborates that dopamine signaling is important in immune cells, which express dopamine receptors (DRD1-DRD5) as well as the molecular machinery for dopamine synthesis and metabolism. Dopamine can regulate inflammatory activity, cell trafficking, and disease pathology, yet the epigenetic mechanisms underlying these effects remain poorly understood. Here, we show that in primary human monocyte-derived macrophages, dopamine increases DNA methylation at the IL-1β proximal promoter in a DNMT-dependent manner, while concurrently upregulating IL-1β gene expression. Dopamine also increases the expression of key epigenetic regulators, including TET2, HDAC2, and HDAC6, suggesting coordinated changes in both DNA methylation and histone modifications that shape inflammatory transcription. Importantly, baseline dopamine receptor expression and donor demographics, including sex and age, influence the magnitude of these epigenetic responses, highlighting inter-individual variability in macrophage sensitivity to dopaminergic signaling. These findings establish dopamine as a modulator of macrophage inflammation via epigenetic remodeling and provide a mechanistic framework for understanding how peripheral immune cells respond to dopaminergic cues. By linking dopamine signaling, epigenetic regulation, and innate immunity, this work identifies potential targets for therapeutic intervention and supports the use of accessible human immune cells to investigate dopaminergic dysregulation in neuroimmunological disorders.

## 1. Introduction

Dopamine is a monoamine neurotransmitter classically associated with reward, motivation, and motor control in the central nervous system (CNS), signaling through five G protein–coupled receptors (DRD1–DRD5) and tightly regulated by the dopamine transporter (DAT). Beyond these canonical mechanisms, dopaminergic activity interfaces with epigenetic pathways that regulate gene expression independently of DNA sequence. These pathways include DNA methylation and post-translational histone modifications, which are mediated by DNA methyltransferases and histone-modifying enzymes, respectively. Through engagement of these epigenetic mechanisms, transient dopamine signals can be translated into stable and enduring changes in transcriptional programs that shape cellular function and behavior over time (Nestler and Lüscher, 2019, Torquet et al., 2018).

Dopamine signaling and epigenetic regulation have a complex, bidirectional interplay. On one hand, dopamine’s canonical actions are shaped by epigenetic mechanisms that regulate the expression of genes involved in dopamine synthesis, signaling, and receptor/transporter function, thereby modulating dopamine availability and influencing cellular responsiveness to dopaminergic cues (Green et al., 2019, Marion-Poll et al., 2022, He et al., 2011, González et al., 2019, Vucetic et al., 2012) (**Table 1**). Given dopamine’s salient role in neurological disease, altered dopaminergic tone can drive persistent epigenetic changes that underlie long-term transcriptional reprogramming in neuropsychiatric and neurodegenerative disorders, including substance use disorders, depression, schizophrenia, and Parkinson’s disease (Ammal Kaidery et al., 2013, Wu et al., 2023, Hillemacher et al., 2019, Muench et al., 2018, Rubino et al., 2020) (**Table 1-3**). Conversely, environmentally or pharmacologically induced changes in dopamine levels can themselves trigger epigenetic modifications, including alterations in DNA methylation and histone marks that reshape chromatin accessibility and downstream gene expression (Penrod et al., 2018, Vilca et al., 2023, Södersten et al., 2014, Authement et al., 2016) (**Table 2**). In addition to these indirect mechanisms, dopamine can directly modify histones through dopaminylation (Lepack et al., 2020), adding another layer of regulation through which dopaminergic signaling shapes cellular plasticity. Collectively, these findings position epigenetic regulation as a critical mediator of dopamine-dependent cellular function and dysfunction.

**Table 1.**
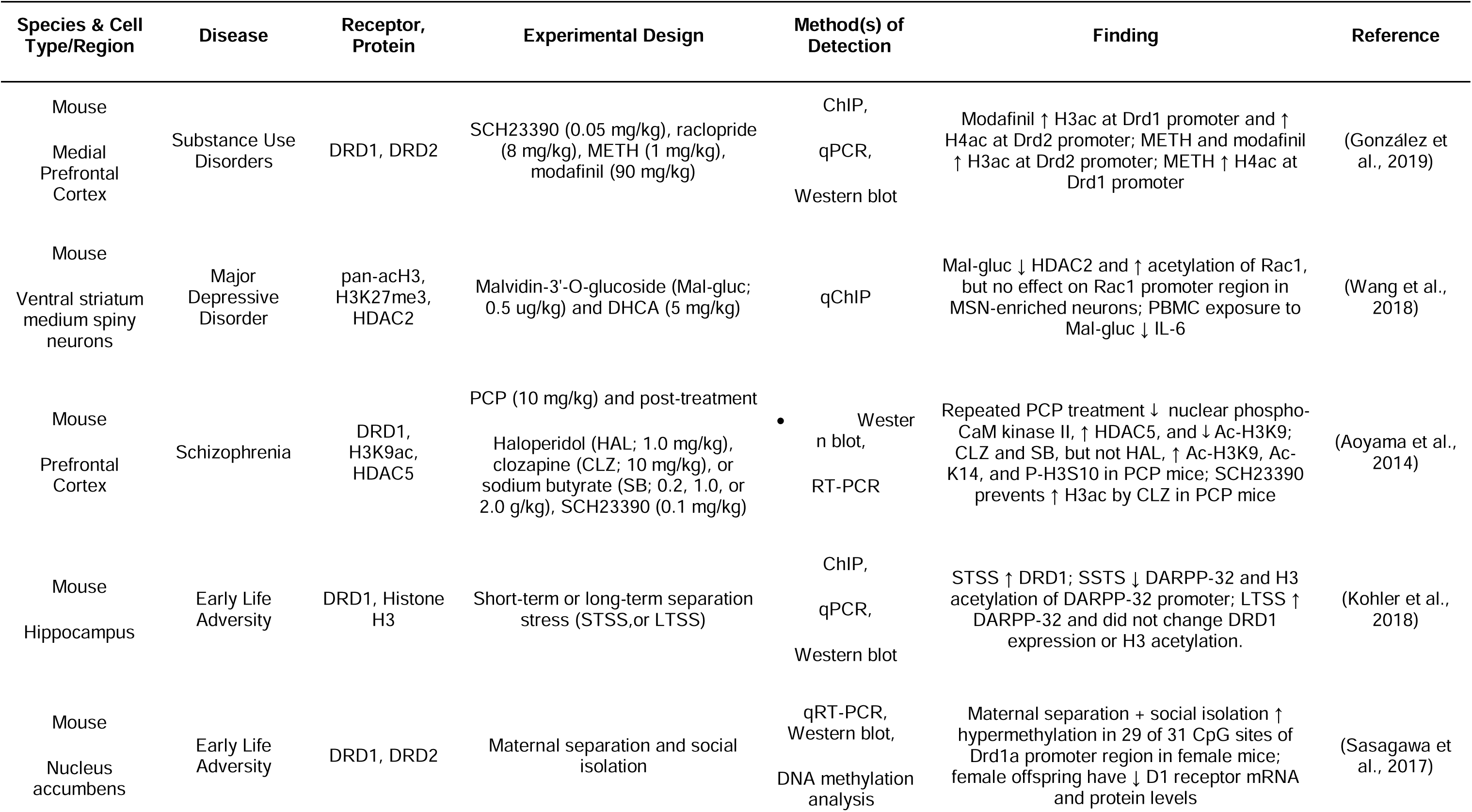
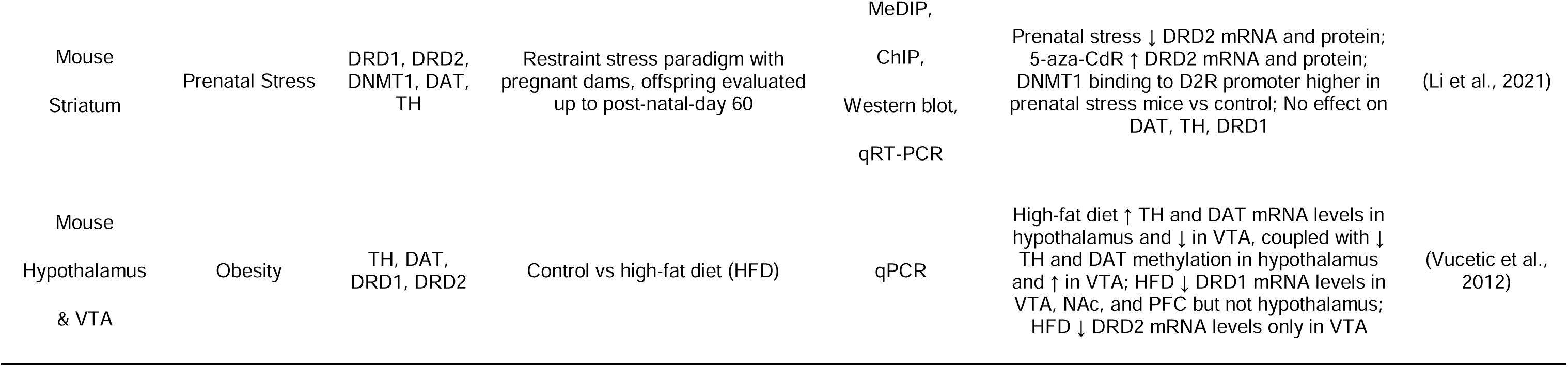
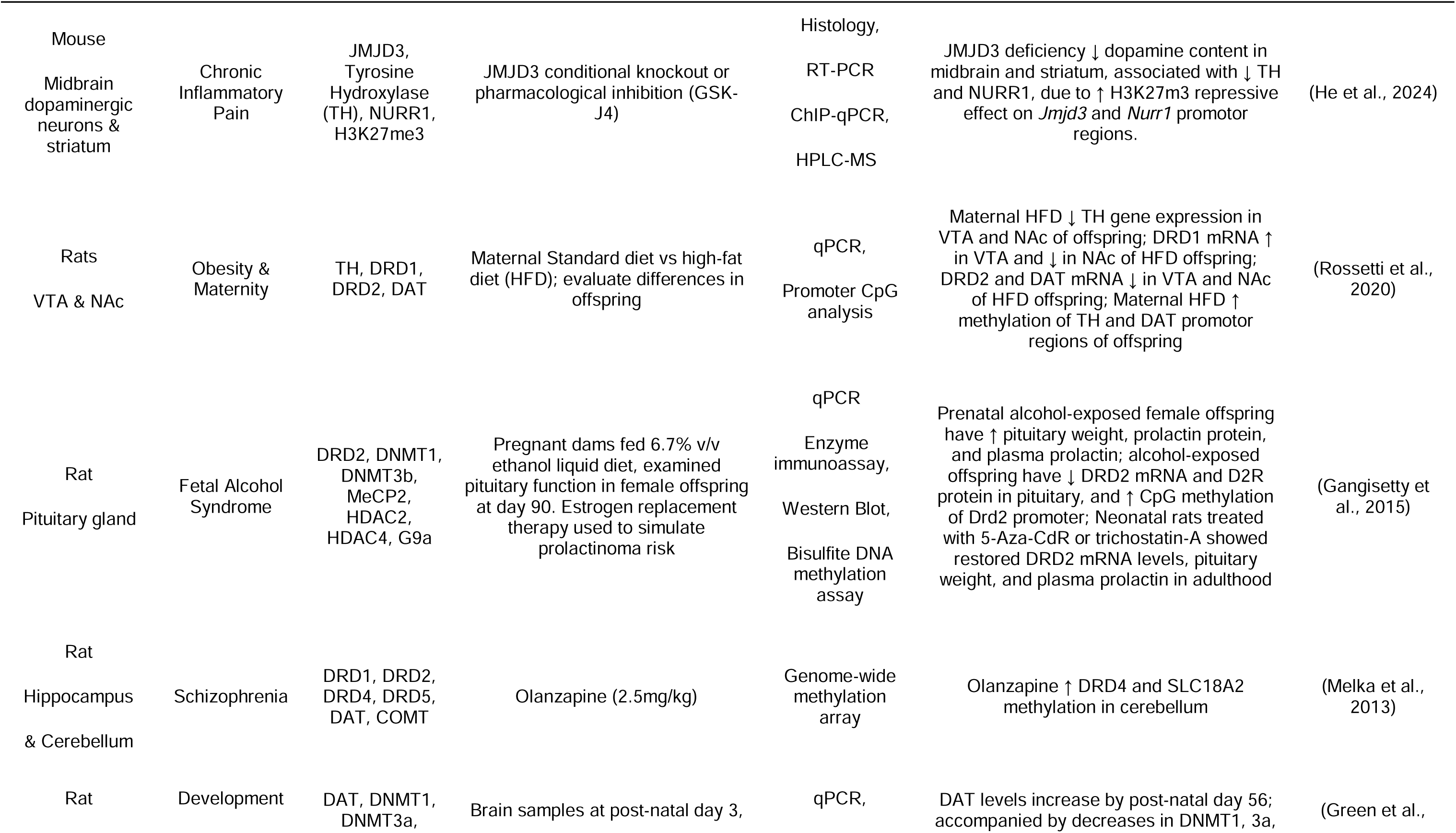
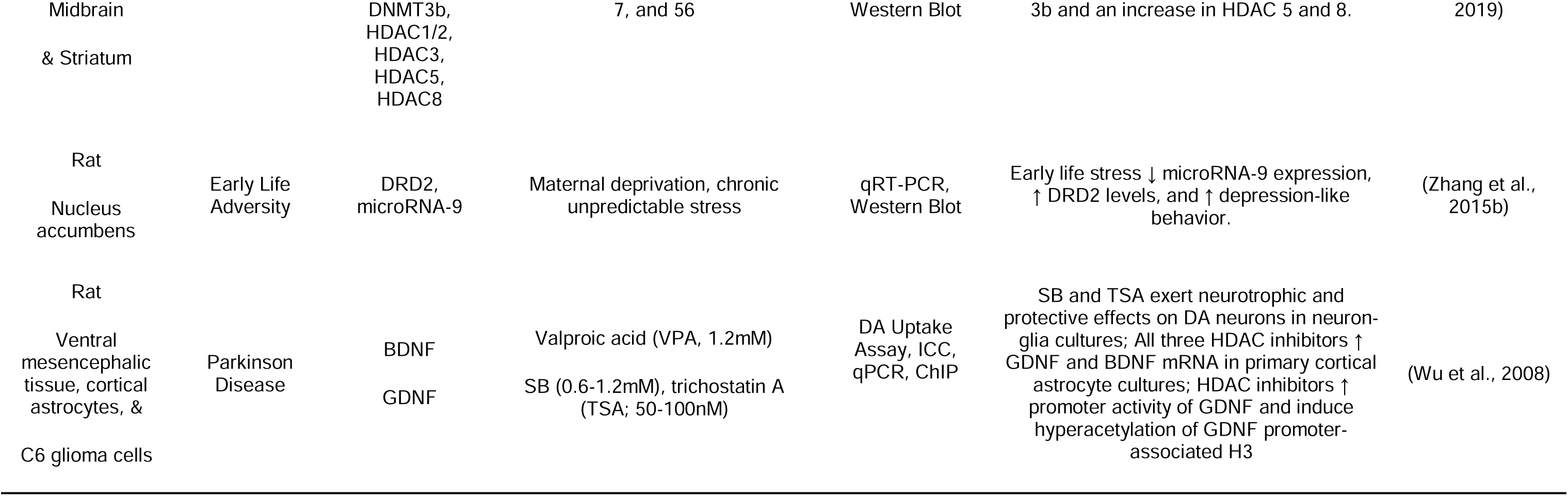
Epigenetic Regulation of Dopaminergic Signaling.

| Species & Cell Type/Region | Disease | Receptor, Protein | Experimental Design | Method(s) of Detection | Finding | Reference |
| --- | --- | --- | --- | --- | --- | --- |
| Mouse<br>Medial Prefrontal Cortex | Substance Use Disorders | DRD1, DRD2 | SCH23390 (0.05 mg/kg), raclopride (8 mg/kg), METH (1 mg/kg), modafinil (90 mg/kg) | ChIP,<br>qPCR,<br>Western blot | Modafinil ↑ H3ac at Drd1 promoter and ↑ H4ac at Drd2 promoter; METH and modafinil ↑ H3ac at Drd2 promoter; METH ↑ H4ac at Drd1 promoter | (González et al., 2019) |
| Mouse<br>Ventral striatum medium spiny neurons | Major Depressive Disorder | pan-acH3, H3K27me3, HDAC2 | Malvidin-3'-O-glucoside (Mal-gluc; 0.5 ug/kg) and DHCA (5 mg/kg) | qChIP | Mal-gluc ↓ HDAC2 and ↑ acetylation of Rac1, but no effect on Rac1 promoter region in MSN-enriched neurons; PBMC exposure to Mal-gluc ↓ IL-6 | (Wang et al., 2018) |
| Mouse<br>Prefrontal Cortex | Schizophrenia | DRD1, H3K9ac, HDAC5 | PCP (10 mg/kg) and post-treatment<br>Haloperidol (HAL; 1.0 mg/kg), clozapine (CLZ; 10 mg/kg), or sodium butyrate (SB; 0.2, 1.0, or 2.0 g/kg), SCH23390 (0.1 mg/kg) | • Western blot,<br>RT-PCR | Repeated PCP treatment ↓ nuclear phospho-CaM kinase II, ↑ HDAC5, and ↓ Ac-H3K9; CLZ and SB, but not HAL, ↑ Ac-H3K9, Ac-K14, and P-H3S10 in PCP mice; SCH23390 prevents ↑ H3ac by CLZ in PCP mice | (Aoyama et al., 2014) |
| Mouse<br>Hippocampus | Early Life Adversity | DRD1, Histone H3 | Short-term or long-term separation stress (STSS, or LTSS) | ChIP,<br>qPCR,<br>Western blot | STSS ↑ DRD1; SSTS ↓ DARPP-32 and H3 acetylation of DARPP-32 promoter; LTSS ↑ DARPP-32 and did not change DRD1 expression or H3 acetylation. | (Kohler et al., 2018) |
| Mouse<br>Nucleus accumbens | Early Life Adversity | DRD1, DRD2 | Maternal separation and social isolation | qRT-PCR,<br>Western blot,<br>DNA methylation analysis | Maternal separation + social isolation ↑ hypermethylation in 29 of 31 CpG sites of Drd1a promoter region in female mice; female offspring have ↓ D1 receptor mRNA and protein levels | (Sasagawa et al., 2017) |
| Dopamine Epigenetically Regulates Macrophage Inflammation |  |  |  |  |  |  |
| Mouse<br>Striatum | Prenatal Stress | DRD1, DRD2,<br>DNMT1, DAT,<br>TH | Restraint stress paradigm with<br>pregnant dams, offspring evaluated<br>up to post-natal-day 60 | MeDIP,<br>ChIP,<br>Western blot,<br>qRT-PCR | Prenatal stress ↓ DRD2 mRNA and protein;<br>5-aza-CdR ↑ DRD2 mRNA and protein;<br>DNMT1 binding to D2R promoter higher in<br>prenatal stress mice vs control; No effect on<br>DAT, TH, DRD1 | (Li et al., 2021) |
| Mouse<br>Hypothalamus<br>& VTA | Obesity | TH, DAT,<br>DRD1, DRD2 | Control vs high-fat diet (HFD) | qPCR | High-fat diet ↑ TH and DAT mRNA levels in<br>hypothalamus and ↓ in VTA, coupled with ↓<br>TH and DAT methylation in hypothalamus<br>and ↑ in VTA; HFD ↓ DRD1 mRNA levels in<br>VTA, NAc, and PFC but not hypothalamus;<br>HFD ↓ DRD2 mRNA levels only in VTA | (Vucetic et al.,<br>2012) |
| Mouse<br>Midbrain dopaminergic neurons & striatum | Chronic Inflammatory Pain | JMJD3, Tyrosine Hydroxylase (TH), NURR1, H3K27me3 | JMJD3 conditional knockout or pharmacological inhibition (GSK-J4) | Histology,<br>RT-PCR<br>ChIP-qPCR,<br>HPLC-MS | JMJD3 deficiency ↓ dopamine content in midbrain and striatum, associated with ↓ TH and NURR1, due to ↑ H3K27m3 repressive effect on *Jmjd3* and *Nurr1* promotor regions. | (He et al., 2024) |
| Rats<br>VTA & NAc | Obesity & Maternity | TH, DRD1, DRD2, DAT | Maternal Standard diet vs high-fat diet (HFD); evaluate differences in offspring | qPCR,<br>Promoter CpG analysis | Maternal HFD ↓ TH gene expression in VTA and NAc of offspring; DRD1 mRNA ↑ in VTA and ↓ in NAc of HFD offspring; DRD2 and DAT mRNA ↓ in VTA and NAc of HFD offspring; Maternal HFD ↑ methylation of TH and DAT promotor regions of offspring | (Rossetti et al., 2020) |
| Rat<br>Pituitary gland | Fetal Alcohol Syndrome | DRD2, DNMT1, DNMT3b, MeCP2, HDAC2, HDAC4, G9a | Pregnant dams fed 6.7% v/v ethanol liquid diet, examined pituitary function in female offspring at day 90. Estrogen replacement therapy used to simulate prolactinoma risk | qPCR<br>Enzyme immunoassay,<br>Western Blot,<br>Bisulfite DNA methylation assay | Prenatal alcohol-exposed female offspring have ↑ pituitary weight, prolactin protein, and plasma prolactin; alcohol-exposed offspring have ↓ DRD2 mRNA and D2R protein in pituitary, and ↑ CpG methylation of *Drd2* promoter; Neonatal rats treated with 5-Aza-CdR or trichostatin-A showed restored DRD2 mRNA levels, pituitary weight, and plasma prolactin in adulthood | (Gangisetty et al., 2015) |
| Rat<br>Hippocampus & Cerebellum | Schizophrenia | DRD1, DRD2, DRD4, DRD5, DAT, COMT | Olanzapine (2.5mg/kg) | Genome-wide methylation array | Olanzapine ↑ DRD4 and SLC18A2 methylation in cerebellum | (Melka et al., 2013) |
| Rat | Development | DAT, DNMT1, DNMT3a, | Brain samples at post-natal day 3, | qPCR, | DAT levels increase by post-natal day 56; accompanied by decreases in DNMT1, 3a, | (Green et al., |
| Dopamine Epigenetically Regulates Macrophage Inflammation |  |  |  |  |  |  |
| Midbrain<br>& Striatum |  | DNMT3b,<br>HDAC1/2,<br>HDAC3,<br>HDAC5,<br>HDAC8 | 7, and 56 | Western Blot | 3b and an increase in HDAC 5 and 8. | 2019) |
| Rat<br>Nucleus<br>accumbens | Early Life<br>Adversity | DRD2,<br>microRNA-9 | Maternal deprivation, chronic<br>unpredictable stress | qRT-PCR,<br>Western Blot | Early life stress ↓ microRNA-9 expression,<br>↑ DRD2 levels, and ↑ depression-like<br>behavior. | (Zhang et al.,<br>2015b) |
| Rat<br>Ventral<br>mesencephalic<br>tissue, cortical<br>astrocytes, &<br>C6 glioma cells | Parkinson<br>Disease | BDNF<br>GDNF | Valproic acid (VPA, 1.2mM)<br>SB (0.6-1.2mM), trichostatin A<br>(TSA; 50-100nM) | DA Uptake<br>Assay, ICC,<br>qPCR, ChIP | SB and TSA exert neurotrophic and<br>protective effects on DA neurons in neuron-<br>glia cultures; All three HDAC inhibitors ↑<br>GDNF and BDNF mRNA in primary cortical<br>astrocyte cultures; HDAC inhibitors ↑<br>promoter activity of GDNF and induce<br>hyperacetylation of GDNF promoter-<br>associated H3 | (Wu et al., 2008) |

**Table 2.**
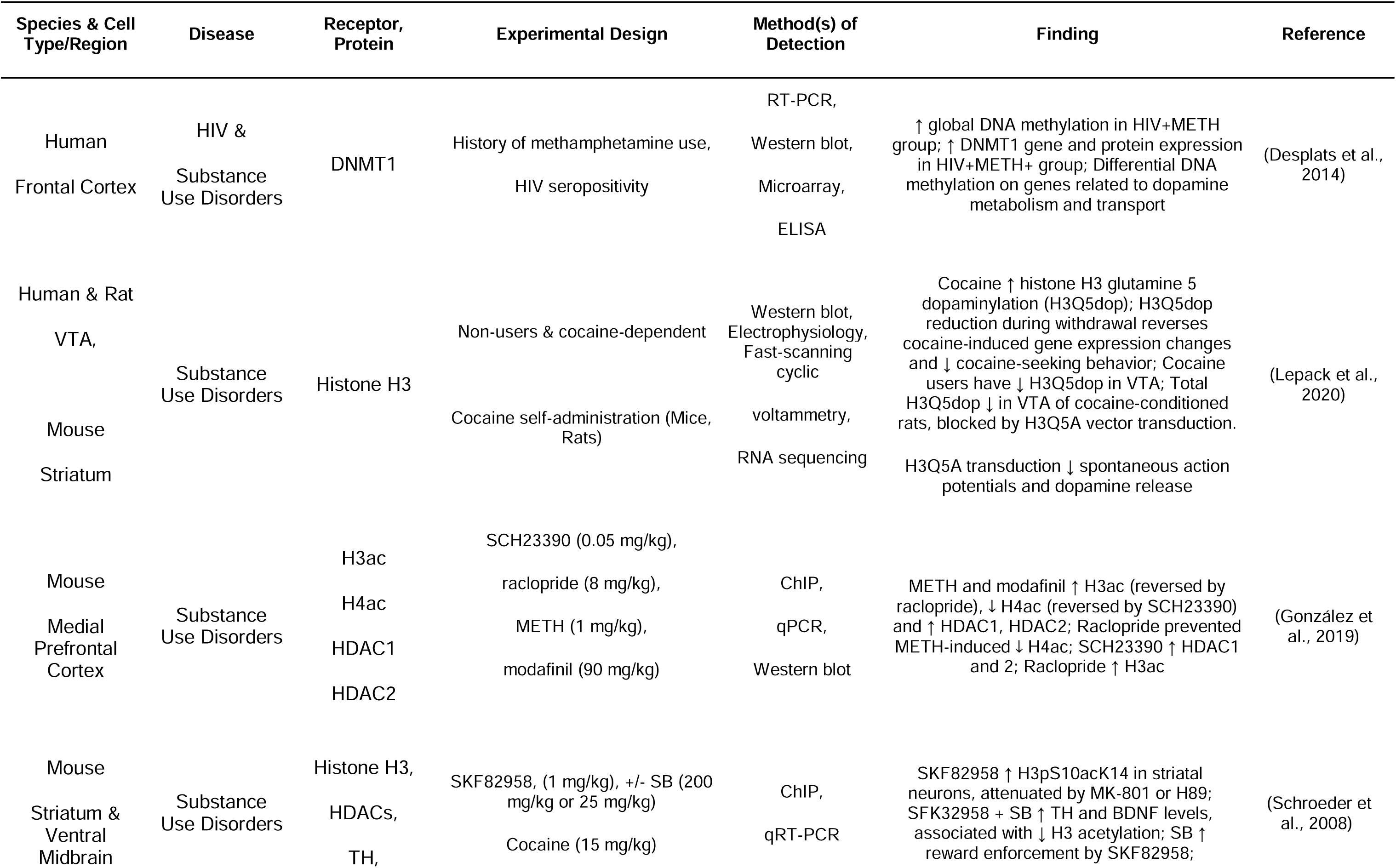
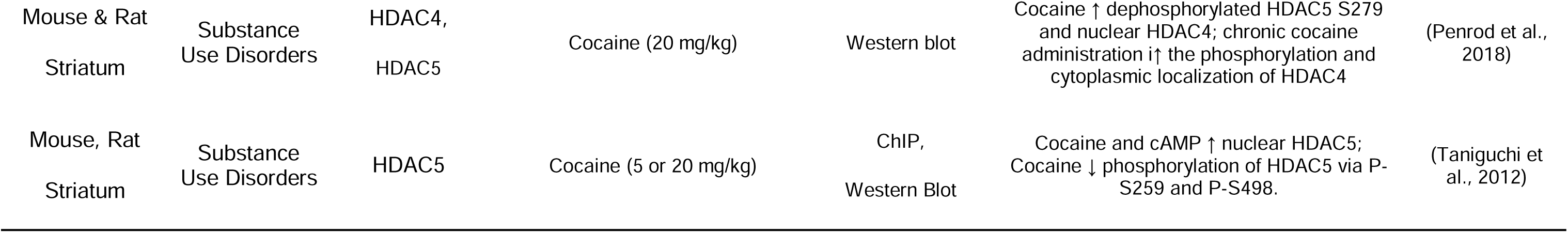
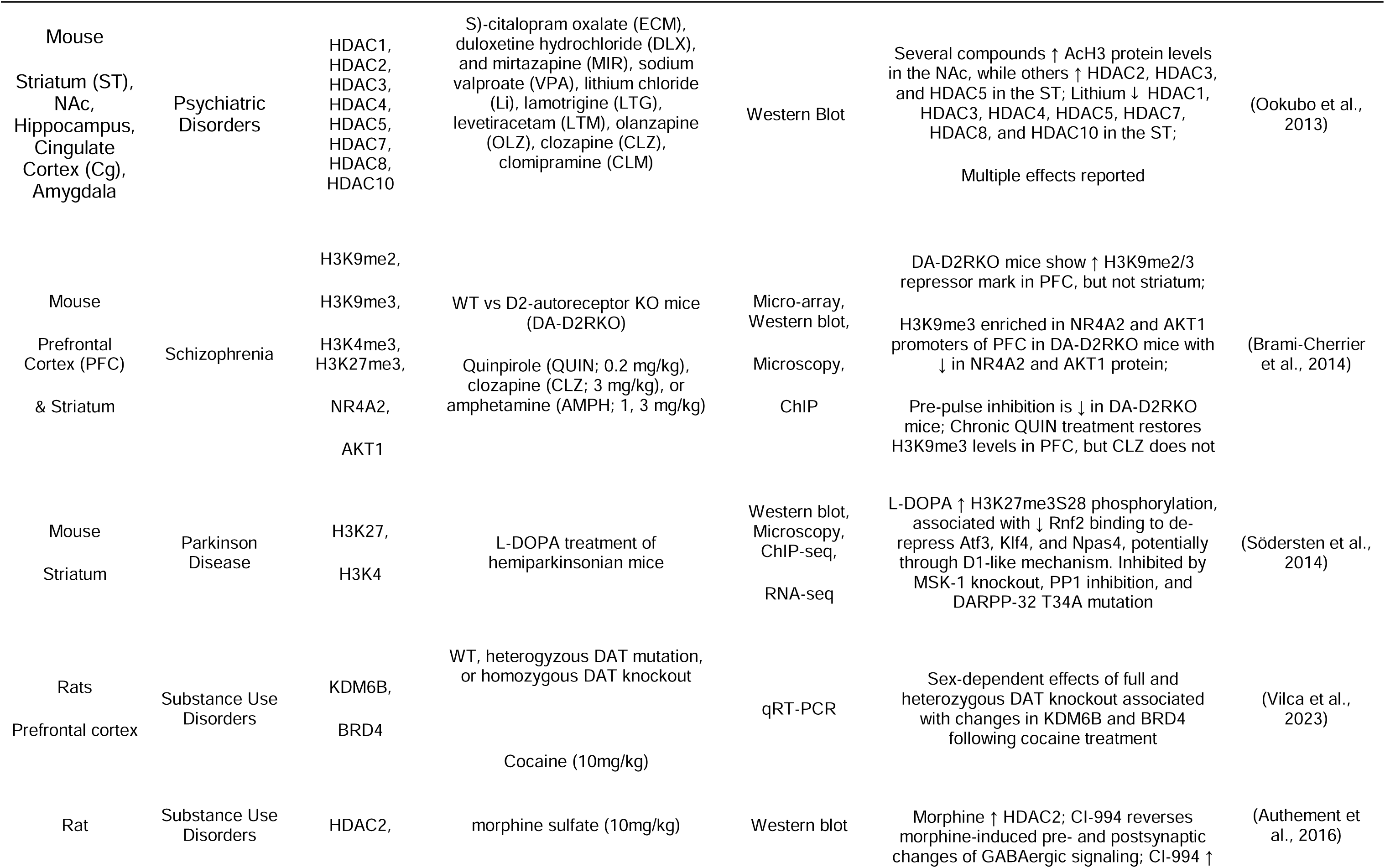
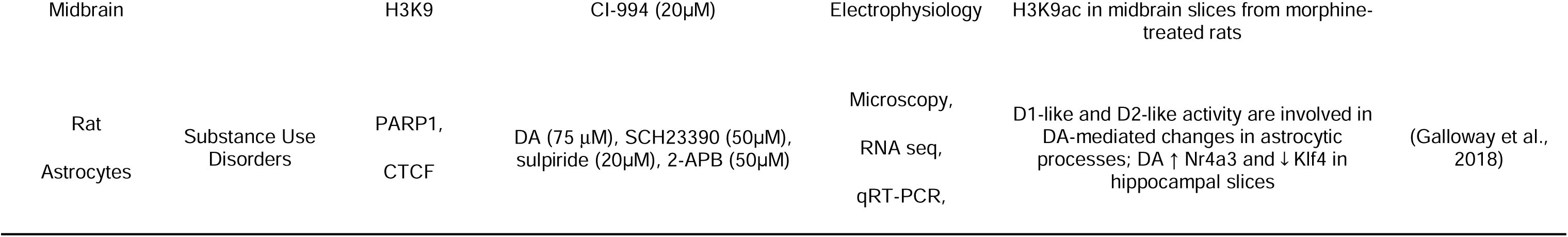
Epigenetic Effects of Dopamine-Modifying Drugs.

**Table 3.**
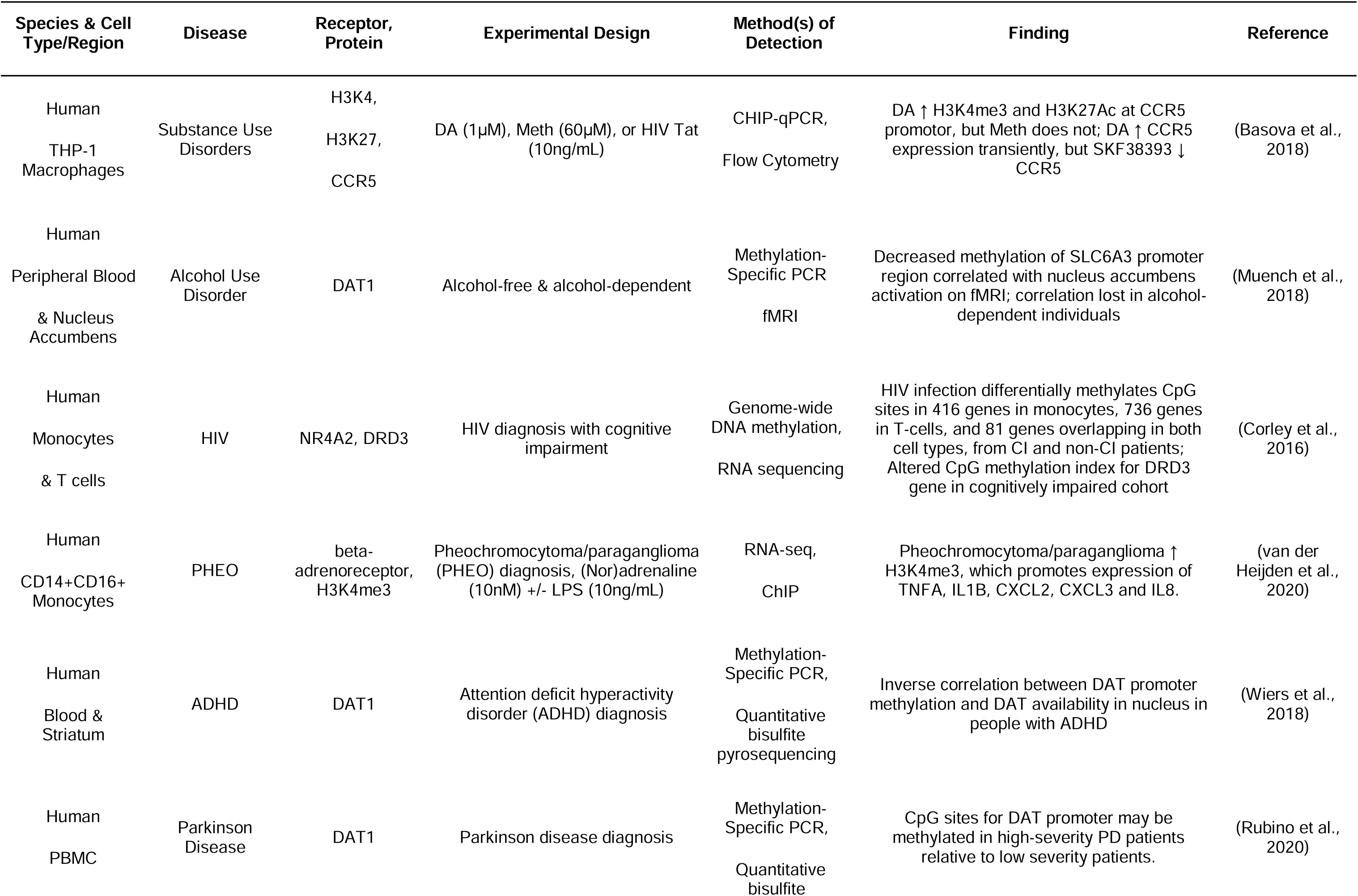
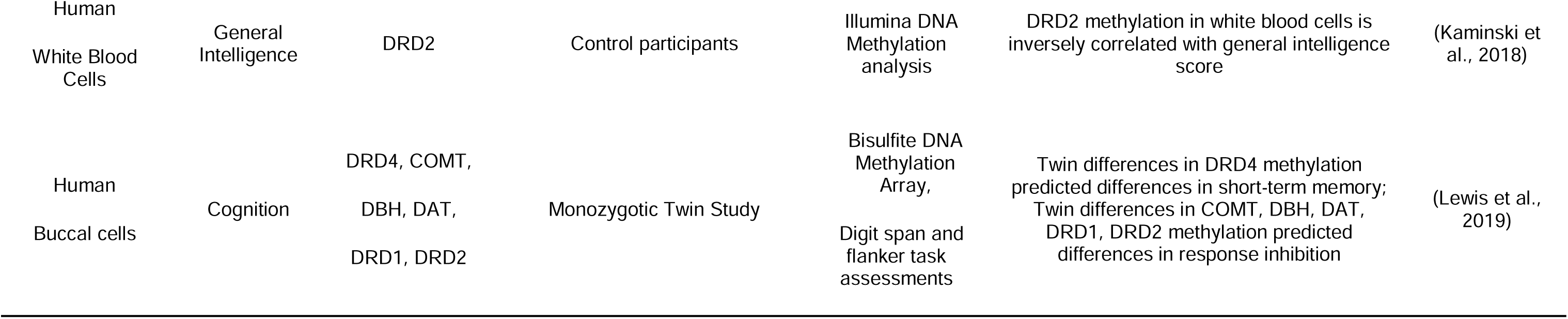
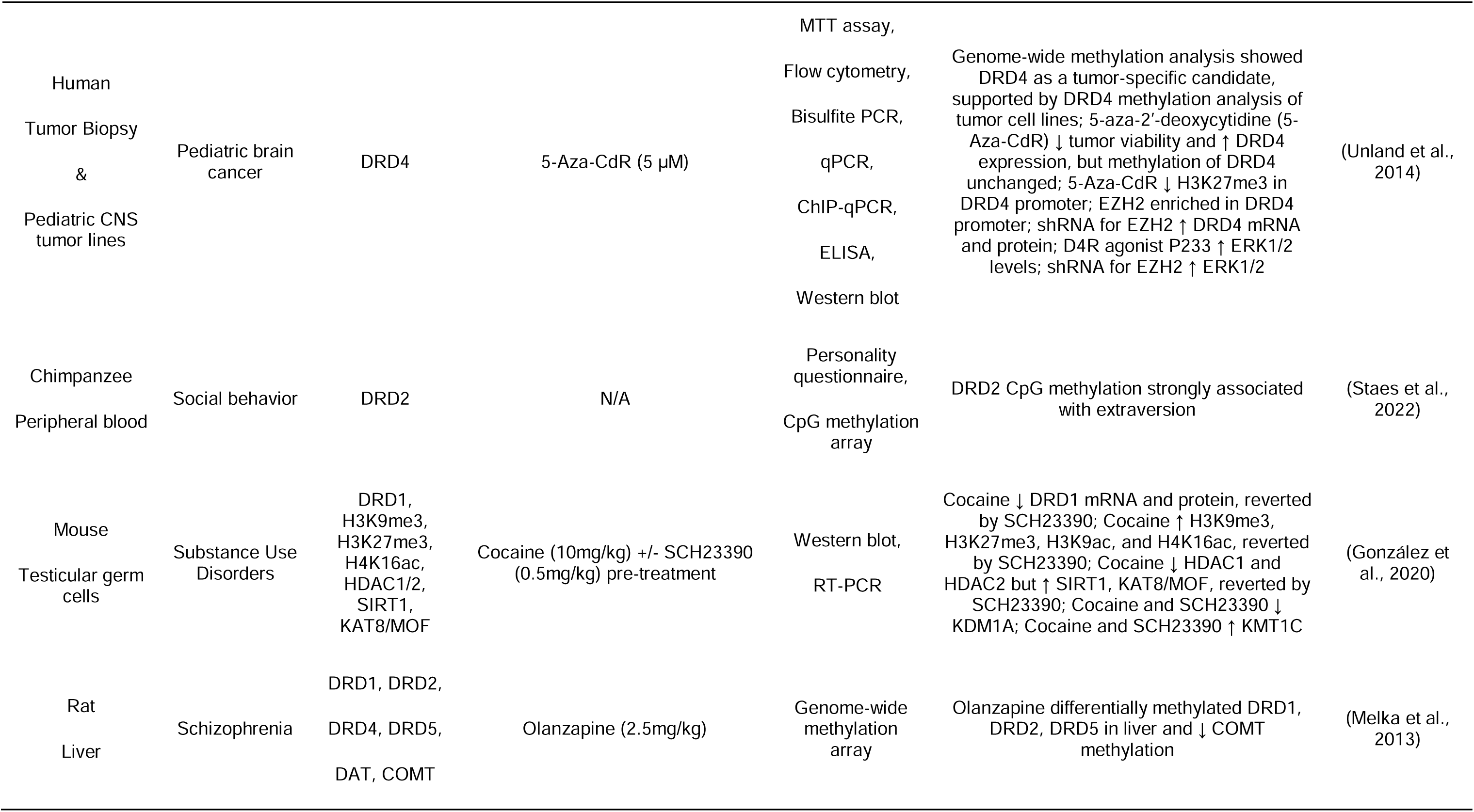
Dopamine-associated regulation of epigenetics in non-neuronal cells.

Dopamine Epigenetically Regulates Macrophage Inflammation
| Species & Cell Type/Region | Disease | Receptor, Protein | Experimental Design | Method(s) of Detection | Finding | Reference |
| --- | --- | --- | --- | --- | --- | --- |
| Human<br>THP-1<br>Macrophages | Substance Use Disorders | H3K4,<br>H3K27,<br>CCR5 | DA (1μM), Meth (60μM), or HIV Tat (10ng/mL) | CHIP-qPCR,<br>Flow Cytometry | DA ↑ H3K4me3 and H3K27Ac at CCR5 promotor, but Meth does not; DA ↑ CCR5 expression transiently, but SKF38393 ↓ CCR5 | (Basova et al., 2018) |
| Human<br>Peripheral Blood<br>& Nucleus<br>Accumbens | Alcohol Use Disorder | DAT1 | Alcohol-free & alcohol-dependent | Methylation-Specific PCR<br>fMRI | Decreased methylation of SLC6A3 promoter region correlated with nucleus accumbens activation on fMRI; correlation lost in alcohol-dependent individuals | (Muench et al., 2018) |
| Human<br>Monocytes<br>& T cells | HIV | NR4A2, DRD3 | HIV diagnosis with cognitive impairment | Genome-wide DNA methylation,<br>RNA sequencing | HIV infection differentially methylates CpG sites in 416 genes in monocytes, 736 genes in T-cells, and 81 genes overlapping in both cell types, from CI and non-CI patients; Altered CpG methylation index for DRD3 gene in cognitively impaired cohort | (Corley et al., 2016) |
| Human<br>CD14+CD16+<br>Monocytes | PHEO | beta-adrenoreceptor,<br>H3K4me3 | Pheochromocytoma/paraganglioma (PHEO) diagnosis, (Nor)adrenaline (10nM) +/- LPS (10ng/mL) | RNA-seq,<br>ChIP | Pheochromocytoma/paraganglioma ↑ H3K4me3, which promotes expression of TNFA, IL1B, CXCL2, CXCL3 and IL8. | (van der Heijden et al., 2020) |
| Human<br>Blood & Striatum | ADHD | DAT1 | Attention deficit hyperactivity disorder (ADHD) diagnosis | Methylation-Specific PCR,<br>Quantitative bisulfite pyrosequencing | Inverse correlation between DAT promoter methylation and DAT availability in nucleus in people with ADHD | (Wiers et al., 2018) |
| Human<br>PBMC | Parkinson Disease | DAT1 | Parkinson disease diagnosis | Methylation-Specific PCR,<br>Quantitative bisulfite | CpG sites for DAT promoter may be methylated in high-severity PD patients relative to low severity patients. | (Rubino et al., 2020) |
| Human<br>White Blood<br>Cells | General<br>Intelligence | DRD2 | Control participants | Illumina DNA<br>Methylation<br>analysis | DRD2 methylation in white blood cells is<br>inversely correlated with general intelligence<br>score | (Kaminski et<br>al., 2018) |
| Human<br>Buccal cells | Cognition | DRD4, COMT,<br>DBH, DAT,<br>DRD1, DRD2 | Monozygotic Twin Study | Bisulfite DNA<br>Methylation<br>Array,<br>Digit span and<br>flanker task<br>assessments | Twin differences in DRD4 methylation<br>predicted differences in short-term memory;<br>Twin differences in COMT, DBH, DAT,<br>DRD1, DRD2 methylation predicted<br>differences in response inhibition | (Lewis et al.,<br>2019) |

Dopamine Epigenetically Regulates Macrophage Inflammation
| Species & Cell Type/Region | Disease | Receptor, Protein | Experimental Design | Method(s) of Detection | Finding | Reference |
| --- | --- | --- | --- | --- | --- | --- |
| Human<br>Tumor Biopsy<br>&<br>Pediatric CNS tumor lines | Pediatric brain cancer | DRD4 | 5-Aza-CdR (5 µM) | MTT assay,<br>Flow cytometry,<br>Bisulfite PCR,<br>qPCR,<br>ChIP-qPCR,<br>ELISA,<br>Western blot | Genome-wide methylation analysis showed DRD4 as a tumor-specific candidate, supported by DRD4 methylation analysis of tumor cell lines; 5-aza-2'-deoxycytidine (5-Aza-CdR) ↓ tumor viability and ↑ DRD4 expression, but methylation of DRD4 unchanged; 5-Aza-CdR ↓ H3K27me3 in DRD4 promoter; EZH2 enriched in DRD4 promoter; shRNA for EZH2 ↑ DRD4 mRNA and protein; D4R agonist P233 ↑ ERK1/2 levels; shRNA for EZH2 ↑ ERK1/2 | (Unland et al., 2014) |
| Chimpanzee<br>Peripheral blood | Social behavior | DRD2 | N/A | Personality questionnaire,<br>CpG methylation array | DRD2 CpG methylation strongly associated with extraversion | (Staes et al., 2022) |
| Mouse<br>Testicular germ cells | Substance Use Disorders | DRD1,<br>H3K9me3,<br>H3K27me3,<br>H4K16ac,<br>HDAC1/2,<br>SIRT1,<br>KAT8/MOF | Cocaine (10mg/kg) +/- SCH23390 (0.5mg/kg) pre-treatment | Western blot,<br>RT-PCR | Cocaine ↓ DRD1 mRNA and protein, reverted by SCH23390; Cocaine ↑ H3K9me3, H3K27me3, H3K9ac, and H4K16ac, reverted by SCH23390; Cocaine ↓ HDAC1 and HDAC2 but ↑ SIRT1, KAT8/MOF, reverted by SCH23390; Cocaine and SCH23390 ↓ KDM1A; Cocaine and SCH23390 ↑ KMT1C | (González et al., 2020) |
| Rat<br>Liver | Schizophrenia | DRD1, DRD2,<br>DRD4, DRD5,<br>DAT, COMT | Olanzapine (2.5mg/kg) | Genome-wide methylation array | Olanzapine differentially methylated DRD1, DRD2, DRD5 in liver and ↓ COMT methylation | (Melka et al., 2013) |

While dopamine-mediated epigenetic regulation has been studied primarily in the context of neuronal dysfunction in neuropsychiatric and neurodegenerative diseases, these disorders are also characterized by chronic inflammation (Channer et al., 2022, Felger and Treadway, 2017, Shi and Yong, 2025). Interactions between dopaminergic signaling and epigenetic regulation occurs across multiple cell types, suggesting substantial crosstalk between peripheral immune regulation and brain function (Basova et al., 2018, Muench et al., 2018, Rubino et al., 2020, Anier et al., 2022) (**Table 3**). Human studies of epigenetic modifications of dopaminergic machinery in peripheral blood have reported associations with cognitive performance and psychiatric symptom severity (Lewis et al., 2019, Suchanecka et al., 2025, Abdolmaleky et al., 2008), and inflammatory stimuli, including infections such as toxoplasmosis or HIV, can induce epigenetic changes that impact dopamine signaling (Xiao et al., 2014, Desplats et al., 2014, Corley et al., 2016). Recently, deficiency of the histone demethylase JMJD3 (KDM6B) was shown to disrupt dopamine biosynthesis in the midbrain and exacerbate chronic inflammatory pain (He et al., 2024), providing direct evidence that epigenetic regulation of dopaminergic pathways can influence inflammatory outcomes. However, existing studies have largely focused on epigenetic regulation of dopaminergic machinery rather than inflammatory gene targets in immune cells, highlighting a critical knowledge gap.

This expanding understanding of dopamine’s involvement in both the brain and the immune system points to a broader role in regulating inflammation throughout the body. Dopamine signaling is not restricted to neuronal cells; it is also present in peripheral tissues, where various immune cell populations express the molecular machinery necessary for dopamine synthesis, metabolism, and signaling. This enables direct dopaminergic regulation of immune function (Channer et al., 2022, Matt and Gaskill, 2019, Thomas Broome et al., 2020, Feng and Lu, 2021). Consistent with this, both endogenous dopamine and dopaminergic drugs have been shown to modulate immune cell activation and inflammatory responses across diverse immunological contexts. Several groups, including our own, have documented dopamine receptor expression and DAT in human myeloid cells (Matt et al., 2021, Gaskill et al., 2012, Gaskill et al., 2014, Yan et al., 2015, Liu et al., 2021, Nickoloff-Bybel et al., 2019, Gopinath et al., 2022, Mackie et al., 2022), supporting that these cells can respond to dopamine. Specifically, we found that dopamine can exert an inflammatory effect on myeloid cells such as primary human monocyte-derived macrophages (MDM) and iPSC-derived microglia, in part by upregulating the NF-kB pathway, increasing NLRP3 priming and IL-1β expression/release (Nolan et al., 2020, Matt et al., 2025). This has been shown to be mediated by levels of dopamine receptor expression, in particular the D1-like receptors (DRD1 and DRD5) (Matt et al., 2025). However, it is not known if dopamine modulates macrophage immune responses through mechanisms that involve epigenetic regulation.

Macrophages are central regulators of innate immune responses and exhibit significant functional plasticity in response to environmental cues. Given their post-mitotic nature and longevity, macrophages face the challenge of maintaining genomic stability while remaining highly adaptable. Epigenetic mechanisms regulate this plasticity by controlling chromatin accessibility at inflammatory gene loci, which in turn facilitates the binding of key transcription factors, such as NF-κB, STAT, and MAPK (Cheray and Joseph, 2018, Chen et al., 2020). This binding is crucial for determining the strength and duration of inflammatory responses in macrophages. In particular, the epigenetic control of genes like IL-1β, a critical mediator of inflammation, plays a pivotal role in the macrophage’s ability to mount an effective immune response. While many studies across various cell types, including macrophages, have examined changes in IL-1β DNA methylation (Matt et al., 2016, Vento-Tormo et al., 2017, Cho et al., 2015, Kirchner et al., 2014), no research has specifically explored the impact of dopamine on IL-1β DNA methylation or other epigenetic modifications in macrophages. Given that macrophages can often be exposed to high dopamine levels and share conserved epigenetic regulatory machinery with neural cells, they represent a biologically relevant and tractable system for studying dopamine-driven epigenetic remodeling. In this study, we investigate the role of epigenetic mechanisms in dopamine-mediated regulation of inflammation in primary human monocyte-derived macrophages. Our findings demonstrate that dopamine increases DNA methylation at the IL-1β proximal promoter in a DNA methyltransferase–dependent manner and increases the expression of key epigenetic regulators, including TET2, HDAC2, and HDAC6. Further, we show that donor demographics and dopamine receptor expression influence these epigenetic changes. Collectively, these findings establish a mechanistic link between dopamine signaling, epigenetic regulation, and macrophage inflammation, providing new insights into how dopaminergic modulation of innate immune responses could contribute to disease pathogenesis.

## 2. Methods

### 2.1 Reagents

RPMI 1640 medium (11875119, Gibco), Fetal Bovine Serum (FBS; MT35010CV, Gibco), Human AB serum (100-512-100, Gemini Bio), HEPES (BP2991), Penicillin/Streptomycin (15140163, Gibco), and sterile dH2O (10977023, Invitrogen) were all purchased from ThermoFisher. Macrophage-colony stimulating factor (M-CSF; 315-02) was obtained from Peprotech. Dopamine hydrochloride (DA; H8502), 5-deoxyazacytidine (dAZA; A3656) and 5-azacytidine (AZA; A2385) were obtained from Sigma-Aldrich. TaqMan Fast Universal Master Mix, and PCR assay probes for DRD1-5 (Hs00265245_s1, Hs00241436_m1, Hs00364455_m1, Hs00609526_m1, Hs00361234_s1), IL1B (Hs01555410_m1), DNMT1 (Hs00945875_m1), DNMT3A (Hs01027166_m1), DNMT3B (Hs00171876_m1), TET2 (Hs00325999_m1), HDAC2 (Hs00231032_m1), HDAC6 (Hs00997427_m1), KDM6B (Hs00996325_g1), KAT2A (Hs00904943_gH), and 18S (4319413E) genes were purchased from Applied Biosystems. Reagents were stored and handled according to manufacturer’s conditions.

Macrophage medium was composed of RPMI 1640 supplemented with 10% heat inactivated FBS, 5% human AB serum, 1% HEPES, 1% Penicillin/Streptomycin, 10ng/mL MCSF. M-CSF lyophilized powder was re-suspended in sterile dH2O at a concentration of 100ug/mL and added to fresh culture medium immediately before use. Dopamine hydrochloride (DA) was resuspended in sterile dH_2_O in the dark at a stock concentration of 10 mM that was then aliquoted and frozen for two months or until use, whichever occurs first. New aliquots were prepared from lyophilized powder every two months. All dopamine preparations and treatments were performed in the dark using 10^-6^M dopamine, with dopamine treatment occurring immediately after thawing the dopamine. dAZA lyophilized powder was resuspended in dH2O to a stock concentration of 10mM and stored at -20C until use. AZA was prepared in DMSO at a stock concentration of 10mM. On the day of the experiment, a fresh vial was thawed and added 1:1000 to culture medium for a final concentration of 1μM.

### 2.2 Cell isolation and culture

Human peripheral blood mononuclear cells (PBMC) were separated from blood obtained from de-identified healthy donors from the New York Blood Center (Long Island City, NY, USA), University of Pennsylvania Human Immunology Core (Philadelphia, PA, USA), and BioIVT (Westbury, NY, USA). Live PBMC counts were estimated using the Nexelcom Cellometer Auto 2000 (Revvity), and 10 million PBMC were used to determine the proportion of monocytes in each donor. Monocyte isolation was performed using the Pan Monocyte Isolation Kit (Miltenyi Biotec, 130-096-537), and the isolated monocytes were counted. According to this result, 950,000 monocytes per well were plated in 6-well plates (Corning) in filter-sterilized macrophage medium (RPMI-1640 supplemented with 10% FBS, 5% human AB serum, 10 mM HEPES, 1% P/S, and 10ng/mL M-CSF). Wells were washed once in plain RPMI medium at three days *in vitro* and replaced with macrophage medium to remove non-adherent PBMCs. Adherent cells matured into monocyte-derived macrophages (MDM) by 6-7 days in culture, as previously described (Matt et al., 2021, Nolan et al., 2020, Nickoloff-Bybel et al., 2019). Limited, de-identified demographic information (age, gender, ethnicity, blood type and cytomegalovirus (CMV) status) was obtained, although all data categories were not available for each donor. All donors (N = 53) were included in analyses of dopamine receptor expression, but incomplete demographic information limited inclusion in some correlation analyses.

### 2.3 Drug Treatments

For dopamine treatments without DNMT inhibitors, mature MDM were treated on day 7 with 1μM dopamine for 3 hours and cell contents were immediately collected in RNA/DNA Lysis Buffer (Zymo Research). For DNMT inhibition experiments, MDM were treated on day 6 (t=0h) with either vehicle (sterile dH2O; 1:1000), 1μM of 5- azacytidine (AZA), or 5-deoxyazatycidine (dAZA) in fresh macrophage medium. After 24h (t=24h), medium was replaced with a second 1μM treatment with vehicle, 1μM 5-AZA, or 1μM 5-dAZA in fresh macrophage medium. After 21h (t=45h from start of treatments), cells were treated with either vehicle or 1μM dopamine for 3 hours. Cell contents were harvested using RNA/DNA Lysis Buffer in the Quick DNA/RNA Miniprep Kit.

### 2.4 Quantitative RT-PCR

Total RNA was extracted from cells using either Trizol (15596026, Invitrogen) or Quick DNA/RNA Miniprep Kit (D7001, Zymo Research), and RNA quantity and purity were determined with the NanoDropOne spectrophotometer (Nanodrop Technologies). RNA (1μg per sample) was used to synthesize cDNA from each donor using the High-Capacity Reverse Transcriptase cDNA Synthesis Kit (4368814, Applied Biosystems). 12.5ng cDNA was used per well, with each gene run in triplicate wells for technical control. TaqMan Universal PCR Master Mix (4304437, Applied Biosystems) was used for the PCR reaction according to manufacturer’s instructions. All dopamine receptor subtypes, epigenetic enzyme genes, and the housekeeping gene 18S were amplified from cDNA by quantitative PCR (qPCR) on a QuantStudio 7 (Applied Biosystems) using gene-specific primers (see reagents section above for primer information).

### 2.5 Methylation-specific Quantitative RT-PCR

Total genomic DNA (gDNA) was extracted using either Trizol (15596026, Invitrogen) or the Quick DNA/RNA Miniprep Kit (D7001, Zymo Research) according to manufacturer’s instructions. gDNA concentration and purity was measured using the NanoDropOne spectrophotometer and was bisulfite-treated using the EZ DNA Methylation Kit (D5001, Zymo Research) according to manufacturer’s instructions. Methylation status of IL1B was assessed via methylation-specific real-time PCR (MSP) using custom primers designed with Methprimer software (http://www. urogene.org/methprimer/) (Li and Dahiya, 2002). Product specificity was determined by melt curve analysis and gel electrophoresis. Primer sets targeted a methylated and unmethylated CG dinucleotide in DNA associated with the promoter region of IL1B (**Table 4**). Unmethylated beta-actin was used as a reference gene as previously published (Liu et al., 2017). Reactions were run using 25ng/mL genomic DNA, 100nM forward and reverse primer per reaction, and iTaq Universal SYBR Green Master Mix (1725121, Bio-Rad), and qPCR was run on a QuantStudio 7 with melt curve analysis for each run. Both high-methylated and low-methylated human genomic DNA (80-8061-HGHM5 and 80-8062-HGUM5 from EpigenDx) were bisulfite treated and used as controls for the primer sets. The high-methylated control amplified for the methylated primer and the low-methylated control amplified for the unmethylated primer. However, amplification for MDMs for the unmethylated primer was not detectable in the majority of samples. This suggests this region is predominantly methylated, and we focused on the methylated primer set.

**Table 4.**
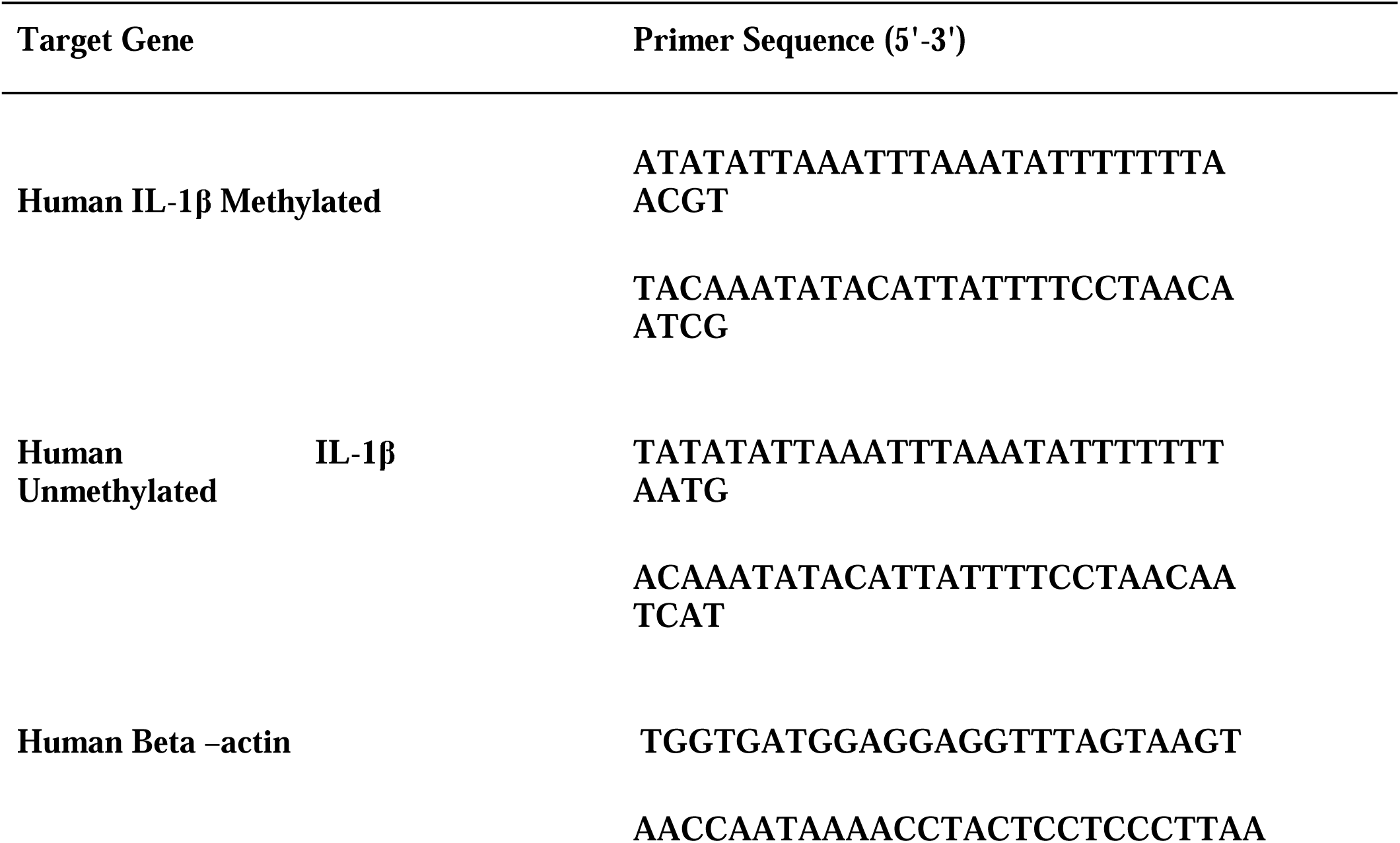
Primers used in MSP experiments.

### 2.6 Statistics

Prior to analysis, all data were normalized to the mean of the vehicle treated condition. Statistical analysis of gene expression data was performed on data normalized to 2^-ΔC^ . To determine the appropriate statistical tests, data sets were evaluated by analysis of skewness and evaluation of normality and lognormality to determine the distribution of the data. Extreme data points presumed to be technical outliers were identified via ROUT test (Q = 0.1%) and removed from analysis. Post-hoc analyses were performed when appropriate. All data analysis was performed using GraphPad Prism 10.2 (Graphpad, La Jolla, CA). p < 0.05 was considered significant.

## 3. Results

### 3.1 DNMT Inhibition Blocks Dopamine-Induced Expression of IL-1β and Dopamine Increases IL-1β DNA Methylation in Primary Human Macrophages

Given the central role of DNA methylation in controlling both signaling of dopamine and inflammation, we investigated whether DNA methylation contributes to dopamine-mediated regulation of IL-1β in primary human monocyte-derived macrophages (MDM). To investigate whether dopamine’s effects on IL-1β in MDM is mediated by DNA methylation, we used two DNMT inhibitors, 5-deoxyazacytidineand (dAZA) and 5-azacytidine (AZA). MDM were pretreated with either DNMT inhibitor (1μM) for 48 hours and then exposed to dopamine for 3 hours at 1μM, which represents a physiologic concentration of dopamine in many tissues throughout the body (Matt and Gaskill, 2019). The 3-hour timepoint was used as we have previously demonstrated dopamine’s effects on inflammation during this window (Matt et al., 2025, Nolan et al., 2020). Cells were then collected for assessment of IL1B mRNA. Pretreatment with dAZA or AZA blocked dopamine’s effects on IL1B (**Figure 1A**, rm 2-way mixed effects model and *post-hoc* with Tukey’s multiple comparisons, dAZA, *p = 0.0260 F (1,7) = 7.913; **Figure 1B**, rm 2-way mixed effects model and *post-hoc* with Tukey’s multiple comparisons, AZA, p = 0.0513 F (1,5) = 6.5), suggesting DNA methylation may play a role in dopamine’s inflammatory effects.

**Figure 1:**
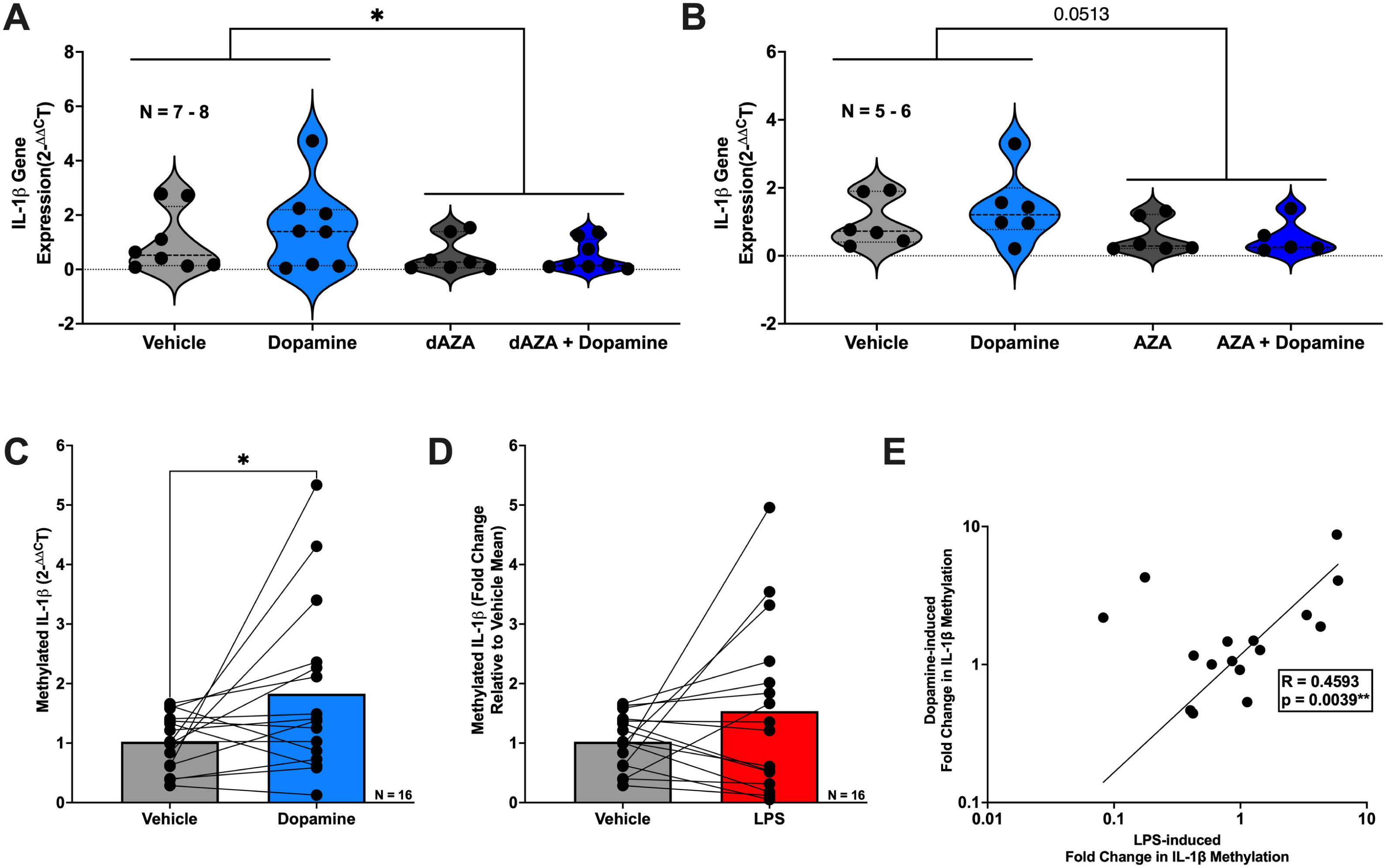
DNMT inhibition blocks dopamine-induced expression of IL-1β and dopamine increases IL-1β DNA methylation in primary human macrophages. Primary human monocyte-derived macrophages (hMDM) from sixteen donors were treated with dopamine (10^-6^M) for 3 hours with or without pre-treatment using hypomethylating agents 5-aza-2’-deoxycytidine (dAZA; 10^-6^M), 5-azacytidine (AZA; 10^-6^M), or vehicle control. RNA and DNA were isolated for analysis of IL-1β gene expression by qPCR and IL-1β DNA methylation, respectively. (A) Dopamine increased IL-1β mRNA expression, an effect inhibited by dAZA (n= 7-8 donors, *p<0.05). (B) AZA pretreatment produced a similar inhibitory effect on dopamine-induced IL-1β expression, trending toward statistical significance (n=5-6, p=0.0513). (C) Dopamine treatment significantly increased methylated IL-1β levels compared to vehicle control (n=16, p<0.05). (D) Lipopolysaccharide (LPS) increased IL-1β DNA methylation; however, this effect did not reach statistical significance in this donor cohort (n=16). (E) LPS-induced changes in IL-1β DNA methylation were significantly correlated with dopamine-induced changes within the same donors, indicating consistent donor-specific responsiveness across stimuli (n=16, **p<0.01).

We then wanted to assess whether specific DNA methylation of IL-1β was affected by dopamine. MDM were treated with 1μM dopamine for 3 hours, and DNA was isolated, bisulfite treated, and examined for IL1B DNA methylation at the proximal promoter, as others have done similarly (Matt et al., 2016, Hashimoto et al., 2009, Tekpli et al., 2013). Dopamine significantly increased IL1B DNA methylation (**Figure 1C**, Vehicle vs Dopamine, n = 16, Wilcoxon test, *p = 0.0335, sum of (+,-) ranks 109, -27), providing a mechanistic basis for the observation that DNMT inhibition blocked dopamine-induced increases in IL1B transcription.

Given that lipopolysaccharide (LPS) is a well-established inducer of IL-1β and is known to influence DNA methylation during myeloid inflammatory responses (Matt et al., 2016, Weinmann et al., 2001, Kruidenier et al., 2012, Jain et al., 2019), we assessed whether LPS (10ng/mL) altered DNA methylation at the IL1B promoter in a manner comparable to dopamine. Although the increase in DNA methylation with LPS did not reach statistical significance (**Figure 1D**, Vehicle vs LPS, n = 16, paired t-test, p = 0.1759, t = 1.421, df = 15), LPS- and dopamine-associated methylation changes in the same MDM donors were positively correlated (**Figure 1E**, n = 16, Pearson r = 0.6777, **p = 0.0039), consistent with the possibility that these stimuli may converge on similar regulatory mechanisms.

### 3.2 Dopamine Increases Expression of Epigenetic Enzymes in Primary Human Macrophages

Given our observation that dopamine increased DNA methylation at the IL1B proximal promoter in a DNMT–dependent manner, we next examined whether dopamine more broadly influences the expression of enzymes that regulate DNA methylation in MDM. We focused on DNMT1, which is primarily responsible for maintenance of DNA methylation patterns during cell division, and DNMT3A/DNMT3B, which mediate de novo DNA methylation, and have all been implicated in regulation of inflammatory gene expression (Yu et al., 2016, Wang et al., 2024, Li et al., 2020). We also examined TET2, a key enzyme involved in active DNA demethylation and resolution of macrophage inflammatory responses (Zhang et al., 2015a, Carrillo-Jimenez et al., 2019, Cull et al., 2017, Sharma et al., 2025, Gao et al., 2023). MDM were treated with 1μM dopamine for 3 hours, and dopamine increased TET2 mRNA (**Figure 2A**, Vehicle vs Dopamine, n = 20, Wilcoxon test, *p = 0.0400, sum of (+,-) ranks 160, -50) while DNMT1, DNMT3A, and DNMT3B mRNA were unchanged (**Figure 2B-D**). This indicates at a global level that dopaminergic signaling may modulate enzymes involved in active DNA demethylation.

**Figure 2:**
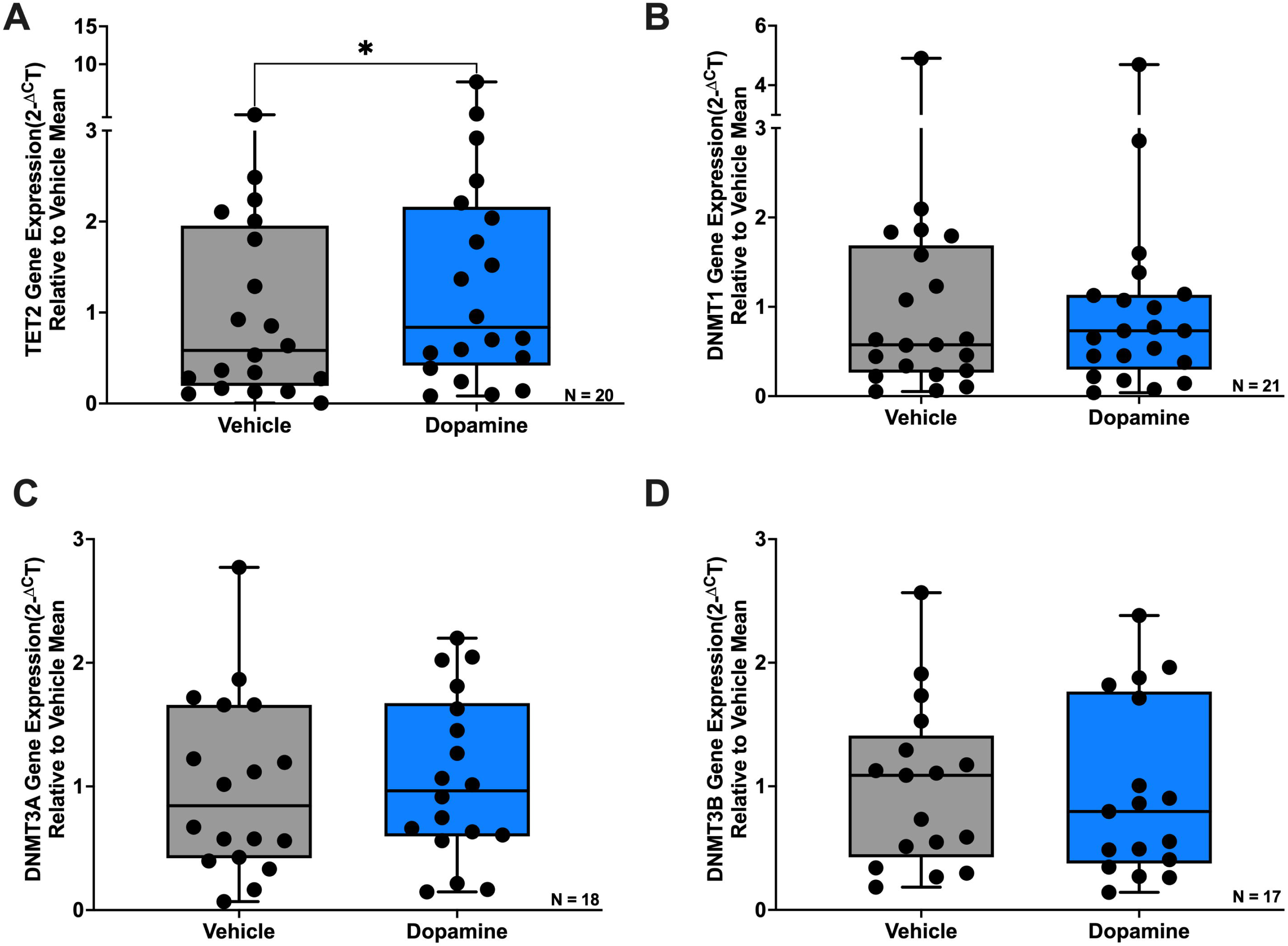
Dopamine selectively increases TET2 expression without altering DNMT expression in primary human macrophages. Primary human monocyte-derived macrophages (hMDM) were treated with dopamine (10^-6^M) for 3 hours, and gene expression was assessed by qPCR. (**A**) Dopamine significantly increased TET2 mRNA expression (n=20 donors, *p<0.05). (**B**) DNMT1 mRNA expression was not significantly altered by dopamine treatment (n=21 donors). (**C**) DNMT3A mRNA expression was not significantly altered by dopamine treatment(n=18 donors). (**D**) DNMT3B mRNA expression was not significantly altered by dopamine treatment (n=17 donors).

In addition to regulating DNA methylation, epigenetic control of inflammatory gene expression in macrophages is mediated by histone modifications, including histone acetylation and methylation, that shape chromatin accessibility (Chen et al., 2020). Given our findings that dopamine modulates enzymes involved in DNA methylation, we next examined whether dopamine also influences the mRNA expression of epigenetic regulators involved in histone modification in MDM. We focused on HDAC2 and HDAC6 due to their established roles in regulating behaviors associated with dopaminergic signaling and substance use disorders (De Carvalho et al., 2021, Fukada et al., 2016, Doke et al., 2021, Yan et al., 2020) as well as inflammation in macrophages through histone deacetylation (Yan et al., 2018, Wu et al., 2019). We also examined KAT2A, a histone acetyltransferase that promotes inflammation by increasing histone acetylation (Zhang et al., 2023b), and KDM6B, a histone demethylase that removes repressive H3K27me3 marks and is involved in macrophage inflammatory signaling (Audu et al., 2022, Johnstone et al., 2021). MDM were treated with 1μM dopamine for 3 hours, and dopamine increased HDAC2 mRNA (**Figure 3A**, Vehicle vs Dopamine, n = 17, Wilcoxon test, *p = 0.0267, sum of (+,-) ranks 123, -30) and HDAC6 mRNA (**Figure 3B**, Vehicle vs Dopamine, n = 14, Wilcoxon test, *p = 0.0353, sum of (+,-) ranks 86, -19), while KAT2A and KDM6B mRNA were unchanged. These results suggest that dopamine’s influence on inflammatory gene expression is not limited to DNA methylation but also extends to histone modifications.

**Figure 3:**
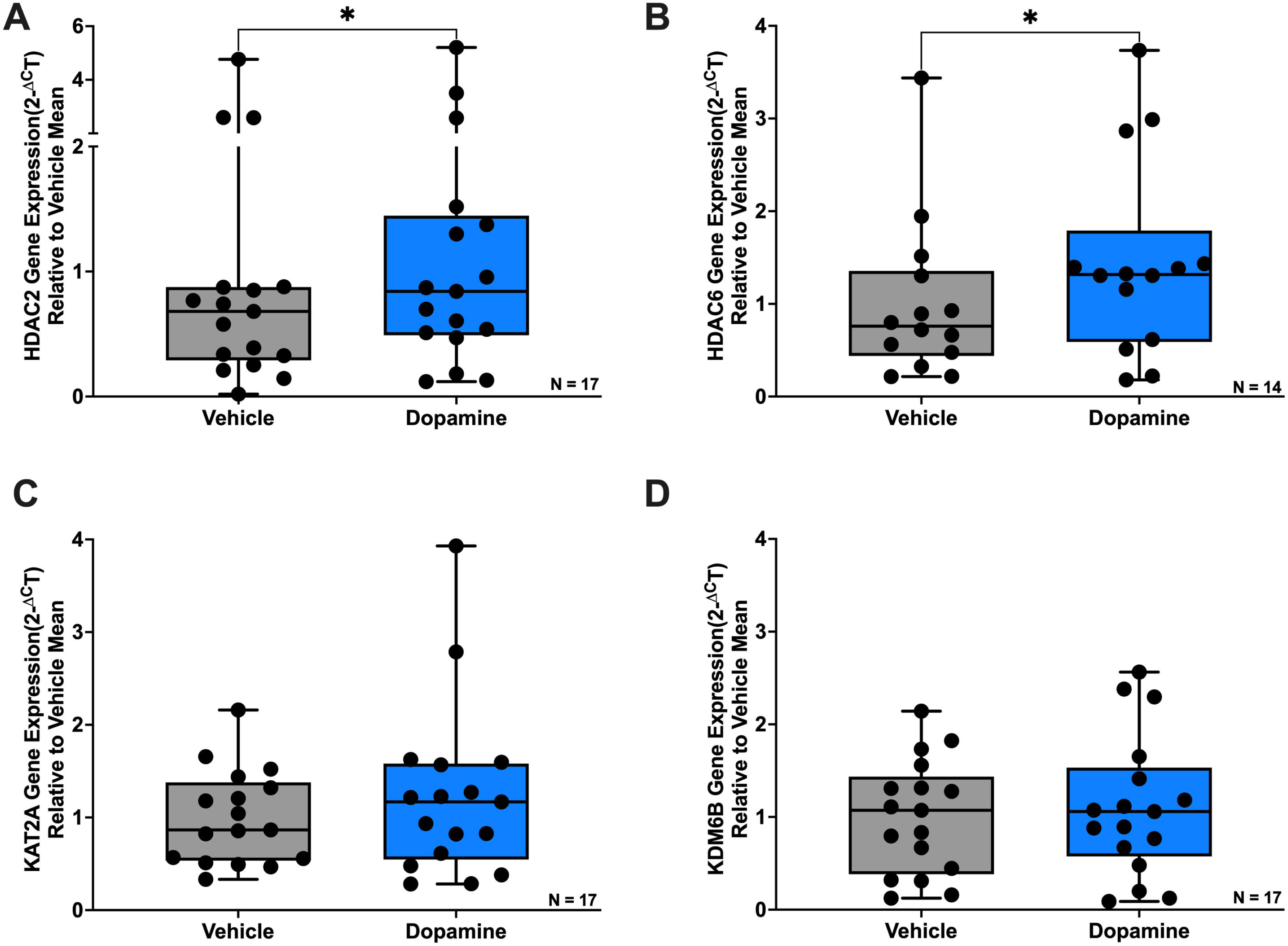
Dopamine increases HDAC2 and HDAC6 expression in primary human macrophages. Primary human monocyte-derived macrophages (hMDM) were treated with dopamine (10^-6^M) for 3 hours, and gene expression was assessed by qPCR. (**A**) Dopamine significantly increased HDAC2 mRNA expression (n=17 donors, *p<0.05). (**B**) Dopamine significantly increased HDAC6 mRNA expression (n=14 donors, *p<0.05). (**C**) KAT2A mRNA expression was not significantly altered by dopamine treatment(n=17 donors). (**D**) KDM6B mRNA expression was not significantly altered by dopamine treatment (n=17 donors).

### 3.3 Sex and Age Influence Dopamine-Induced Changes in Epigenetic Enzymes in Human Macrophages

Having established that dopamine modulates IL-1β DNA methylation and the expression of enzymes involved in DNA methylation and histone modifications in macrophages (MDM), we next considered the known influence of biological sex and age on both dopaminergic signaling and immune responses. We therefore evaluated whether these demographic factors contributed to variability in dopamine-mediated epigenetic regulation. Our analyses revealed that the magnitude of dopamine-induced epigenetic changes varied by sex (males versus females) and age (donors under 40 versus over 40). While no significant differences were observed in IL-1β DNA methylation (data not shown), males exhibited a 2.78-fold increase in DNA methylation, compared to a 1.26-fold increase in females. Similarly, donors under 40 had a 2.34-fold increase, while donors over 40 showed a 1.52-fold increase.

For TET2, dopamine significantly increased its expression in males, but this effect was not observed in females (**Figure 4A**, Males Vehicle vs Dopamine, n = 10, Wilcoxon test, *p = 0.0273, sum of (+,-) ranks 49, -6). Interestingly, for DNMT3A and DNMT3B, while overall dopamine did not significantly alter their expression, there were trends suggesting dopamine decreased their expression in males (**Figure 4C**, Males Vehicle vs Dopamine, n = 9, paired t-test, p = 0.0712, t = 2.4079, df = 8; **Figure 4D**, Males Vehicle vs Dopamine, n = 11, paired t-test, p = 0.0823, t = 1.931, df = 10). Regarding age, only a trend toward increased TET2 expression in donors under 40 was observed (**Figure 4E**, Under 40 Vehicle vs Dopamine, n = 10, Wilcoxon test, p = 0.0645, sum of (+,-) ranks 46, -9), with no significant changes in donors over 40.

**Figure 4:**
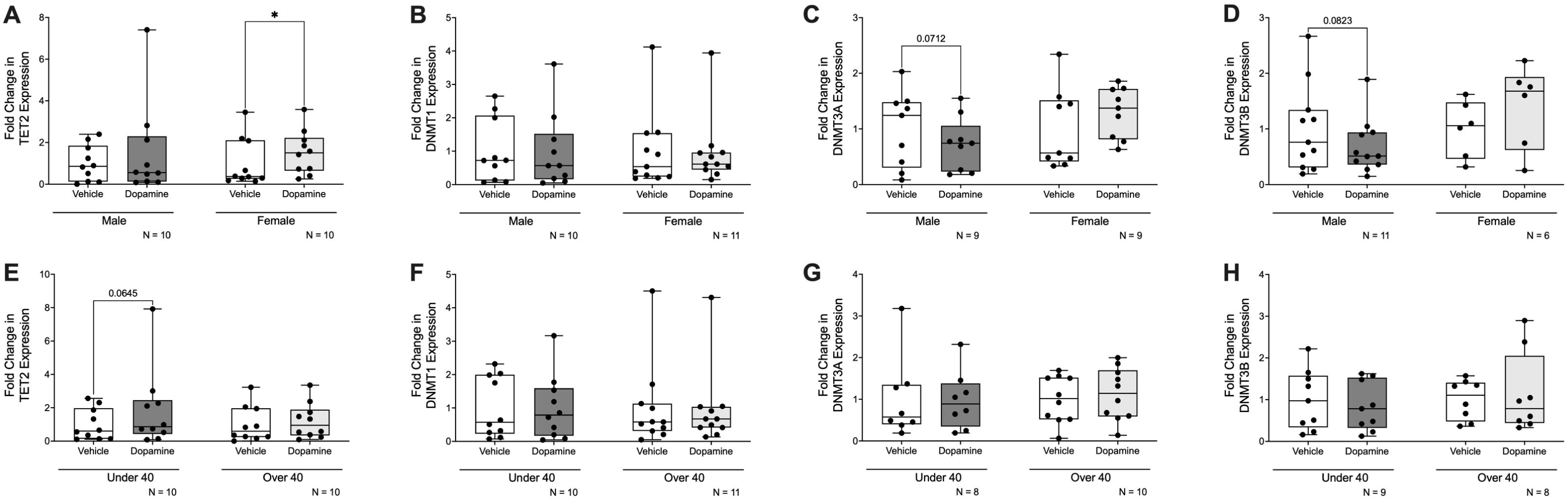
Sex and age influence dopamine-associated regulation of enzymes involved in DNA methylation in primary human macrophages. Data from Figure 2 were stratified by donor sex and age to assess potential sources of inter-donor variability. (**A**) Dopamine significantly increased TET2 mRNA expression in hMDM from female donors but not male donors (n=10 per group, *p<0.05). (**B**) Dopamine did not significantly alter DNMT1 gene expression in either male (n=10) or female (n=11) hMDM. (**C**) Dopamine treatment showed a decreasing trend in DNMT3A expression in male hMDM (n=9; p=0.0712) and a non-significant increase in female hMDM (n=9). (**D**) Dopamine treatment showed a decreasing trend in DNMT3B expression in male hMDM (n=11; p=0.0823) and a non-significant increase in female hMDM (n=6). (**E**) Dopamine treatment showed an increasing trend in TET2 gene expression in donors under 40 years of age (n=10, p=0.0645), but not donors aged 40 years or older (n=10). (**F-H**) No age-associated differences were observed for dopamine-induced changes in (F) DNMT1, (G) DNMT3A, or (H) DNMT3B expression.

For HDAC2, dopamine significantly increased its expression in females, but this effect was not observed in males (**Figure 5A**, Females Vehicle vs Dopamine, n = 10, Wilcoxon test, *p = 0.0488, sum of (+,-) ranks 47, -8). In contrast, dopamine showed a trend toward increased HDAC6 expression in males, but this was not observed in females (**Figure 5B**, Males Vehicle vs Dopamine, n = 9, Wilcoxon test, p = 0.0742, sum of (+,-) ranks 38, -7). For age, there was only a trend toward increased HDAC2 expression in donors under 40 (**Figure 5E**, Under 40 Vehicle vs Dopamine, n = 8, Wilcoxon test, p = 0.0547, sum of (+,-) ranks 32, -4), with no significant change in donors over 40.

**Figure 5:**
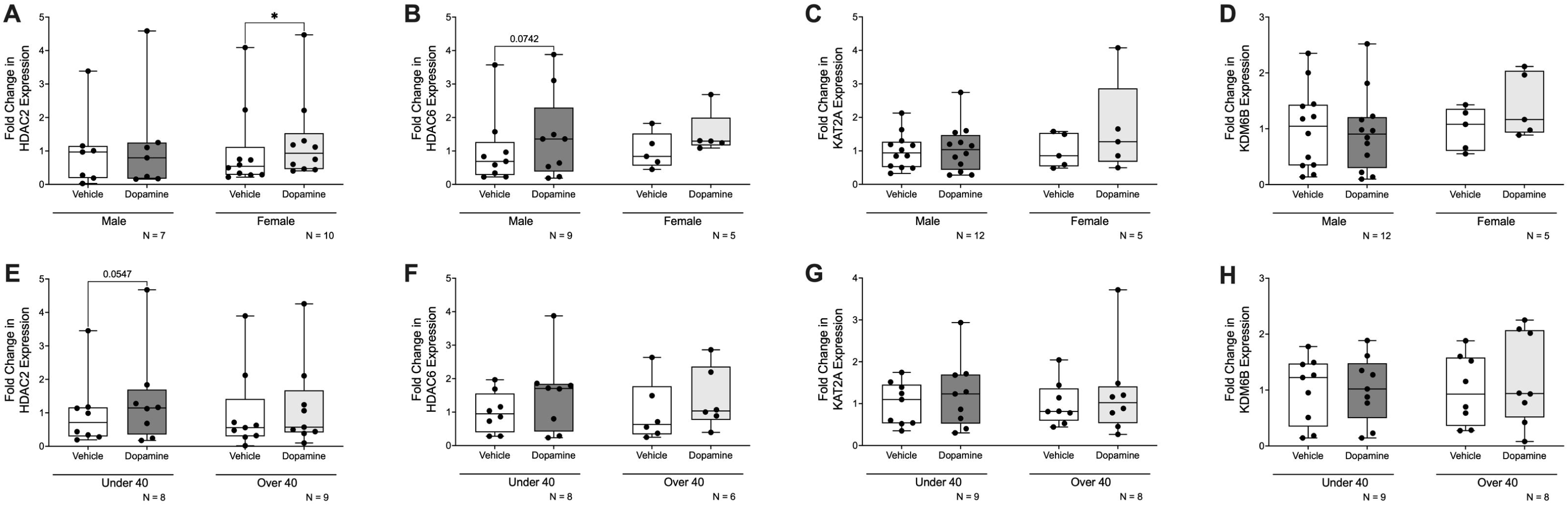
Sex and age influence dopamine-associated regulation of enzymes involved in histone acetylation and methylation in primary human macrophages. Data from Figure 3 were stratified by donor sex and age to assess potential sources of inter-donor variability. (**A**) Dopamine significantly increased HDAC2 mRNA expression in hMDMs from female donors (n=10, *p<0.05) but not male donors (n=7). (**B**) Dopamine treatment showed an increasing trend in HDAC6 expression in male hMDMs (n=9; p=0.0742) and a non-significant increase in female hMDMs (n=5). Dopamine treatment did not reveal sex-associated differences in (**C**) KAT2A or (**D**) KDM6B gene expression (n=17 donors each). (**E**) Dopamine treatment showed an increasing trend in HDAC2 expression in donors under 40 years of age (n=8, p=0.0547), but not in donors aged 40 years or older (n=9). No age-associated differences were observed for dopamine-induced changes in (**F**) HDAC6 (n=14), (**G**) KAT2A (n=17), or (**H**) KDM6B (n=17) expression.

Together, these findings highlight the potential role of demographic factors in shaping macrophage epigenetic responsiveness to dopamine. Even when no overall change in enzyme expression was observed, demographic breakdowns revealed subtle effects that may have been masked in the aggregate analysis. These results suggest that sex and age should be considered as important factors when studying dopamine’s impact on immune function and epigenetic regulation in macrophages.

### 3.4 Epigenetic Enzyme Expression Predominantly Correlates with D1-like Dopamine Receptor Expression in Human Macrophages

We next investigated whether variation in dopamine receptor mRNA expression (DRD1-DRD5) was associated with differences in IL-1β DNA methylation and epigenetic enzyme levels. While no significant correlations were observed between dopamine receptor levels and IL-1β DNA methylation (data not shown), baseline MDM expression of several epigenetic enzymes showed robust positive associations with dopamine receptor expression. Notably, the strongest correlations were observed with D1-like receptors, DRD1 and DRD5. Specifically, TET2, DNMT1, and DNMT3A exhibited significant positive correlations with DRD1, while DNMT3B showed a similar trend (**Figure 6A-D**, TET2 vs DRD1, n = 32, Spearman r = 0.4265, *p = 0.0149; DNMT1 vs DRD1, n = 29, Spearman r = 0.5030, **p = 0.0054; DNMT3A vs DRD1, n =30, Spearman r = 0.5874, ***p = 0.0006; DNMT3B vs DRD1, n =30, Spearman r = 0.3761, p = 0.0532). Comparable positive correlations were also observed for TET2, DNMT1, and DNMT3A with DRD5 (**Figure 6E-G**, TET2 vs DRD5, n = 32, Spearman r = 0.3310, p = 0.0642; DNMT1 vs DRD5, n = 29, Spearman r = 0.3842, *p = 0.0396; DNMT3A vs DRD5, n =30, Spearman r = 0.4287, *p = 0.0181).

**Figure 6:**
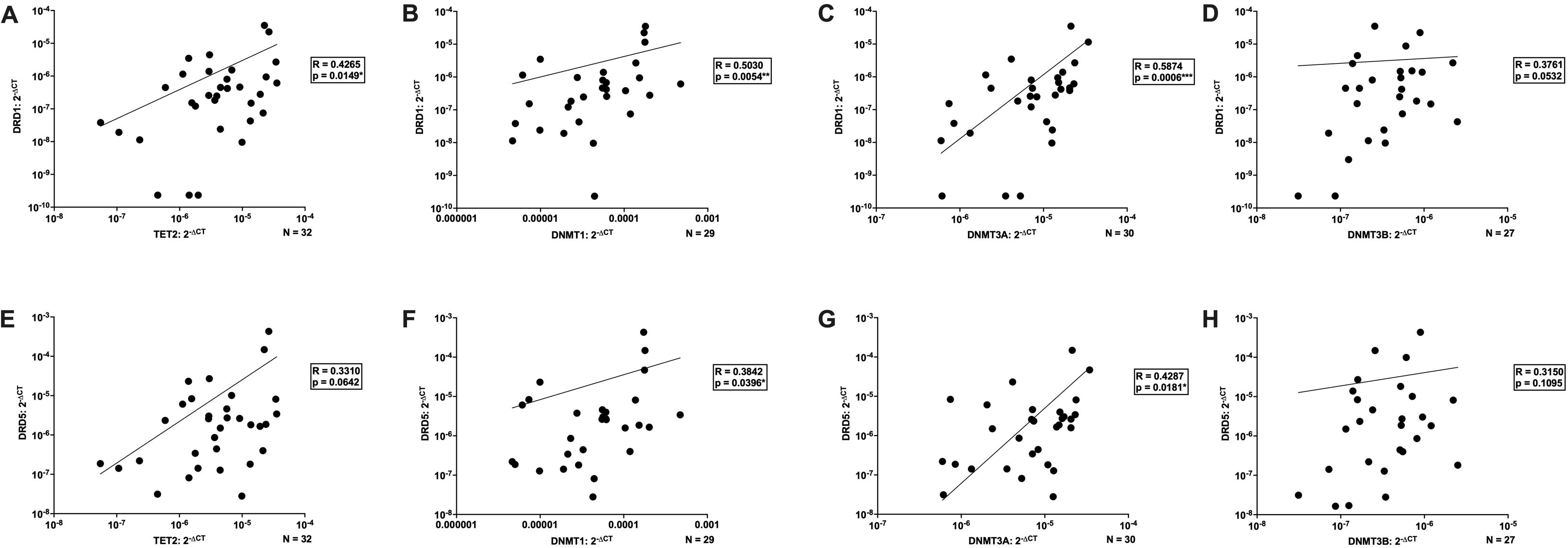
Baseline D1-like receptor expression correlates with enzymes involved in DNA methylation in primary human macrophages. Donors from Figure 2 were combined with an independent archival cohort to assess correlations between dopamine receptor expression and epigenetic enzyme expression by qPCR. DRD1 expression was significantly correlated with (**A**) TET2 (n=32, *p<0.05), (**B**) DNMT1 (n=27, **p<0.01), and (**C**) DNMT3A (n=30, ***p<0.001), but not with (**D**) DNMT3B expression (n=27, p=0.0532). DRD5 expression was not correlated with (**E**) TET2 (n=32, p=0.0642) or (**H**) DNMT3B (n=27), but was significantly correlated with (**F**) DNMT1 (n=29, *p<0.05) and (**G**) DNMT3A (n=30, *p<0.05).

For the histone modifying enzymes, HDAC2, HDAC6, KAT2A, and KDM6B all displayed significant positive correlations with DRD1 (**Figure 7A-D**, HDAC2 vs DRD1, n = 28, Spearman r = 0.4902, **p = 0.0081; HDAC6 vs DRD1, n = 25, Spearman r = 0.5735, **p = 0.0027; KAT2A vs DRD1, n = 28, Spearman r = 0.4803, **p = 0.0097; KDM6B vs DRD1, n = 28, Spearman r = 0.5887, ***p = 0.001). Similarly, these enzymes also showed positive correlations with DRD5 (**Figure 7E-H**, HDAC2 vs DRD5, n = 28, Spearman r = 0.4368, *p = 0.0201; HDAC6 vs DRD5, n = 25, Spearman r = 0.5138, **p = 0.0086; KAT2A vs DRD5, n = 28, Spearman r = 0.3437, p = 0.0733; KDM6B vs DRD5, n = 28, Spearman r = 0.4423, *p = 0.0184).

**Figure 7:**
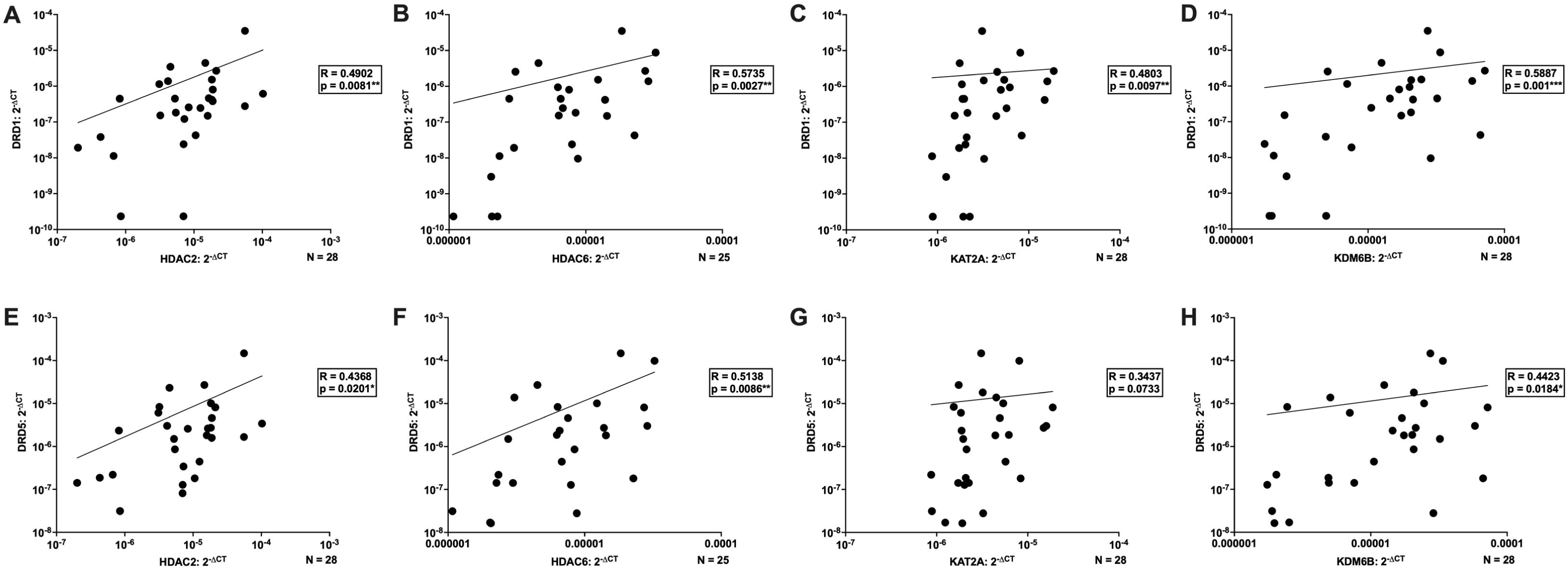
Baseline D1-like dopamine receptor expression correlates with involved in histone acetylation and methylation in primary human macrophages. Donors from Figure 3 were combined with an independent archival cohort to assess correlations between dopamine receptor expression and epigenetic enzyme expression by qPCR. DRD1 expression was significantly correlated with (**A**) HDAC2 (n=28, **p<0.01), (**B**) HDAC6 (n=25, **p<0.01), (**C**) KAT2A (n=28, **p<0.01), and (**D**) KDM6B expression (n=28, ***p<0.001). DRD5 expression was significantly correlated with (**E**) HDAC2 (n=28, *p<0.01), (**F**) HDAC6 (n=25, *p<0.01), and (**H**) KDM6B (n=28, *p<0.05), but not with (**G**) KAT2A expression (n=28, p=0.0733).

In contrast, few associations were observed for D2-like receptors. Only DNMT1 exhibited a significant positive correlation with DRD3 (DNMT1 vs DRD3, n = 29, Spearman r = 0.4394, *p = 0.0171, data not shown), and DNMT3B showed a trend toward a negative correlation with DRD4 (DNMT3B vs DRD4, n = 27, Spearman r = -0.3557, p = 0.0686, data not shown).

Overall, these findings indicate that epigenetic enzyme expression in macrophages is predominantly associated with D1-like receptor levels. This pattern is consistent with our previous observations that dopamine-mediated regulation of macrophage pro-inflammatory signaling is largely driven by D1-like receptors (Nolan et al., 2018, Matt et al., 2025), highlighting a potential mechanistic link between D1-like receptor expression and epigenetic regulation in immune cells. Notably, variability in baseline dopamine receptor expression across donors may establish individual signaling setpoints that constrain the magnitude and direction of epigenetic responses to dopamine.

## 4. Discussion

In this study, we demonstrate that dopamine increases inflammatory gene expression in primary human macrophages through epigenetic mechanisms. Dopamine exposure altered IL-1β expression in a DNA methylation-dependent manner, as pharmacological inhibition of DNMT activity abolished dopamine-induced transcriptional changes. In parallel, dopamine increased the expression of multiple epigenetic enzymes, including TET2, HDAC2, and HDAC6, which are already well-established as key regulators of inflammatory gene expression in myeloid cells (Cull et al., 2017, Wu et al., 2019, Yan et al., 2018, Iwamoto et al., 2020). Together, these results suggest dopamine can act as a novel modulator of inflammatory gene expression through mechanisms previously implicated in classical macrophage activation.

The requirement for DNMT activity implicates DNA methylation as a key mediator of dopamine-induced inflammatory responses. While promoter methylation is typically associated with transcriptional repression, it can have context-dependent effects. Indeed, although dopamine increases IL-1β promoter DNA methylation, it concurrently increased IL-1β expression, and we have previously shown it can increase IL-1β production (Matt et al., 2025, Nolan et al., 2020). This apparent paradox may reflect the dynamic nature of protein–DNA interactions: methylation can block transcription factor binding directly or via recruitment of methyl-CpG-binding proteins, but it can also create novel binding sites or alter transcription factor sequence specificity (Zhu et al., 2016). Future studies could directly assess dopamine’s impact on transcription factor binding at key inflammatory loci such as IL-1β. In addition to NF-κB, other factors should be explored such as AP-1, which is regulated by DRD1 signaling (Kashihara et al., 1999) and whose binding sites are enriched in many genes upregulated by substances of misuse (Zhang et al., 2004), or negative regulators like SP1 (Hashimoto et al., 2013). Understanding which transcription factors are recruited or blocked in response to dopamine will be critical for linking epigenetic remodeling to functional changes in inflammatory gene expression.

Although enzyme expression does not directly equate to enzymatic activity, the concurrent dopamine-induced changes in HDAC2 and HDAC6 expression further suggest that dopamine engages dynamic chromatin remodeling processes, potentially coordinating DNA methylation with histone modifications to fine-tune inflammatory gene expression. Given the highly variable kinetics of epigenetic enzyme activity across the genome (Ginno et al., 2020), dopamine-induced modulation of epigenetic enzymes may produce disproportionate, gene-specific effects on inflammatory responses. These dynamic, dopamine-driven epigenetic changes raise the intriguing possibility that dopamine may contribute to trained immunity–like phenomena in macrophages. Trained immunity describes the persistence of epigenetic and metabolic reprogramming following transient stimuli, resulting in altered inflammatory responsiveness upon subsequent challenges (Netea et al., 2016, Capriotti and Klase, 2024). Catecholamines have been shown to induce trained immunity in monocytes (van der Heijden et al., 2020), and our findings suggest that dopamine may similarly trigger lasting epigenetic events capable of modulating macrophage inflammatory setpoints. By analogy to activity-dependent DNA methylation in neurons, which underlies learning and memory (Bayraktar et al., 2020, Day et al., 2013), dopamine-mediated epigenetic remodeling in myeloid cells could establish long-term changes in responsiveness to inflammatory cues, particularly in tissues with high dopaminergic tone. While trained immunity was not directly assessed in this study, these results suggest a mechanism by which dopamine may shape the capacity of innate immune cells to respond to future stimuli, with potential implications for neuroinflammation and dopaminergic disorders.

Interestingly, we observed that dopamine-induced epigenetic changes varied with donor sex and age. Males and younger donors (<40 years) generally exhibited greater modulation of epigenetic enzyme expression, including DNMTs and HDACs, compared to females and older donors. These findings suggest that biological variables can influence the sensitivity of macrophages to dopaminergic signaling. Consistent with this, we also observed that baseline dopamine receptor expression correlates with epigenetic enzyme levels, particularly for D1-like receptors (DRD1 and DRD5). Several enzymes, including TET2, DNMT1, DNMT3A, HDAC2, HDAC6, KAT2A, and KDM6B, showed positive associations with DRD1 and, to a lesser extent, DRD5. Although these results are limited by cohort size, they suggest that inter-individual variation due to donor demographics or dopamine receptor expression may contribute to heterogeneity in dopamine-mediated epigenetic and inflammatory responses. Importantly, these findings also underscore that dopamine signaling in macrophages is fundamentally distinct from canonical neuronal dopamine signaling (Nickoloff-Bybel et al., 2019, Liu et al., 2021). Non-neuronal cells express different receptor repertoires and engage alternative downstream pathways, which likely shape the context-specific epigenetic consequences of dopamine exposure. In this light, the receptor–enzyme correlations observed in macrophages provide mechanistic insight into how peripheral immune cells may interpret dopaminergic cues differently from neurons.

Epigenetic studies of dopaminergic dysregulation have traditionally focused on brain regions implicated in substance use disorders and neuropsychiatric disease. However, limited access to human brain tissue poses a major barrier to mechanistic investigations. Emerging evidence suggests that peripheral immune cells may serve as informative surrogates: stimulant drugs such as cocaine induce coordinated changes in DNA methylation machinery in both brain regions, including the nucleus accumbens, and in peripheral blood leukocytes, with correlated alterations in DNMT and TET activity (Anier et al., 2022, Anier et al., 2018). Our findings extend this framework by demonstrating that dopamine directly reshapes epigenetic programs in human macrophages, supporting a role for peripheral immune cells as both reporters and contributors to dopaminergic and inflammatory processes. Larger, well-characterized cohorts and genome-wide epigenomic profiling of patient-derived immune cells could further uncover dopamine-associated epigenetic signatures across neuropsychiatric and neurodegenerative conditions.

At the same time, accessible human models of the brain are critical for dissecting CNS-specific mechanisms, making brain organoids a valuable direction for future studies. With regards to substance use disorders, drug-induced dopaminergic transmission is known to alter epigenetic pathways, but it remains unclear whether these effects occur directly or via intermediate dopaminergic signaling. Dopamine agonists alone can modulate histone acetylation, DNA promoter methylation, and HDAC activity (Schroeder et al., 2008, González et al., 2019), yet the epigenetic impact of individual substances of misuse in the absence of intrinsic dopaminergic tone remains largely unexplored. Emerging non-animal models, such as iPSC-derived neural systems, including brain organoids cultured with CNS myeloid cells like microglia (Sun et al., 2024, Kofman et al., 2022, Zhang et al., 2023a), offer a powerful platform to interrogate human-specific molecular responses to addictive substances with or without endogenous dopamine. Coupled with high-resolution single-cell analyses, these approaches provide the opportunity to dissect nuanced effects of neurotransmitter dysfunction on epigenetic regulation, effectively bridging insights from peripheral immune cells to central nervous system processes.

In conclusion, our study identifies dopamine as an epigenetic regulator of inflammatory gene expression in human macrophages, highlighting a previously underappreciated role in modulating innate immune function. By reshaping myeloid epigenetic programs, dopamine may contribute to the pathophysiology of neuropsychiatric and neurodegenerative disorders marked by both dopaminergic dysfunction and immune activation. Characterizing these myeloid-specific epigenetic mechanisms opens avenues for precise therapeutic strategies: coupled with emerging gene-editing and epigenetic-modifying technologies (Marei, 2025), this knowledge could guide the development of targeted interventions, inform patient stratification, and reveal potential contraindications, positioning dopamine-sensitive immune pathways as a critical nexus for translational research.

## Author Contributions

YA, PJG, and SMM contributed to the design and conception of the study. YA, PJG, and SMM designed and analyzed the experiments, and YA, MR, JM, SM, KM, and SMM performed the experiments. PJG and SMM helped supervise the project. YA and SMM performed the statistical analyses and wrote the manuscript. All authors contributed to manuscript revision, read, and approved the final submitted version, and SMM was responsible for the final approval of the submitted version.

## Acknowledgments

We would like to state our tremendous appreciation for all the individuals who donated the biological materials used in these studies. We would also like to thank all the members of the Matt and Gaskill labs for their contributions to this work.

## References

1. Abdolmaleky, H. M., Smith, C. L., Zhou, J. R. & Thiagalingam, S. 2008. Epigenetic alterations of the dopaminergic system in major psychiatric disorders. Methods Mol Biol, 448, 187–212.

2. Ammal Kaidery, N., arannum, S. & Thomas, B. 2013. Epigenetic landscape of Parkinson’s disease: emerging role in disease mechanisms and therapeutic modalities. Neurotherapeutics, 10, 698–708.

3. Anier, K., Somelar, K., Jaako, K., Alttoa, M., Sikk, K., Kokassaar, R., Kisand, K. & Kalda, A. 2022. Psychostimulant-induced aberrant Dna methylation in an in vitro model of human peripheral blood mononuclear cells. Clinical Epigenetics, 14, 89.

4. Anier, K., Urb, M., Kipper, K., Herodes, K., Timmusk, T., Zharkovsky, A. & Kalda, A. 2018. Cocaine-induced epigenetic DNA modification in mouse addiction-specific and non-specific tissues. Neuropharmacology, 139, 13–25.

5. Aoyama, Y., Mouri, A., Toriumi, K., Koseki, T., Narusawa, S., Ikawa, N., Mamiya, T., Nagai, T., Yamada, K. & Nabeshima, T. 2014. Clozapine ameliorates epigenetic and behavioral abnormalities induced by phencyclidine through activation of dopamine D1 receptor. Int J Neuropsychopharmacol, 17, 723–37.

6. Audu, C. O., Melvin, W. J., Joshi, A. D., Wolf, S. J., Moon, J. Y., Davis, F. M., Barrett, E. C., Mangum, K. D., Deng, H., Xing, X., Wasikowski, R., Tsoi, L. C., Sharma, S. B., Bauer, T. M., Shadiow, J., Corriere, M. A., Obi, A. T., Kunkel, S. L., Levi, B., Moore, B. B., Gudjonsson, J. E., Smith, A. M. & Gallagher, K. A. 2022. Macrophage-specific inhibition of the histone demethylase JMJD3 decreases Sting and pathologic inflammation in diabetic wound repair. Cell Mol Immunol, 19, 1251–1262.

7. Authement, M. E., Langlois, L. D., Kassis, H., Gouty, S., Dacher, M., Shepard, R. D., Cox, B. M. & Nugent, F. S. 2016. Morphine-induced synaptic plasticity in the Vta is reversed by Hdac inhibition. J Neurophysiol, 116, 1093–103.

8. Basova, L., Najera, J. A., Bortell, N., Wang, D., Moya, R., Lindsey, A., Semenova, S., Ellis, R. J. & Marcondes, M. C. G. 2018. Dopamine and its receptors play a role in the modulation of CCR5 expression in innate immune cells following exposure to Methamphetamine: Implications to Hiv infection. PLos One, 13, e0199861.

9. Bayraktar, G., Yuanxiang, P., Confettura, A. D., Gomes, G. M., Raza, S. A., Stork, O., Tajima, S., Suetake, I., Karpova, A., Yildirim, F. & Kreutz, M. R. 2020. Synaptic control of Dna methylation involves activity-dependent degradation of DNMT3A1 in the nucleus. Neuropsychopharmacology, 45, 2120–2130.

10. Brami-Cherrier, K., Anzalone, A., Ramos, M., Forne, I., Macciardi, F., Imhof, A. & Borrelli, E. 2014. Epigenetic reprogramming of cortical neurons through alteration of dopaminergic circuits. Mol Psychiatry, 19, 1193–200.

11. Capriotti, Z. & Klase, Z. 2024. Innate immune memory in chronic Hiv and Hiv-associated neurocognitive disorders (Hand): potential mechanisms and clinical implications. J Neurovirol, 30, 451–476.

12. Carrillo-Jimenez, A., Deniz, Ö., Niklison-Chirou, M. V., Ruiz, R., Bezerra-SALOMão, K., Stratoulias, V., Amouroux, R., Yip, P. K., Vilalta, A., Cheray, M., Scott-Egerton, A. M., Rivas, E., Tayara, K., GARCía-DOMínguez, I., Garcia-Revilla, J., Fernandez-Martin, J. C., Espinosa-Oliva, A. M., Shen, X., St George-Hyslop, P., Brown, G. C., Hajkova, P., Joseph, B., Venero, J. L., Branco, M. R. & Burguillos, M. A. 2019. TET2 Regulates the Neuroinflammatory Response in Microglia. Cell Rep, 29, 697–713.e8.

13. Channer, B., Matt, S. M., Nickoloff-Bybel, E. A., Pappa, V., Agarwal, Y., Wickman, J. & Gaskill, P. J. 2022. Dopamine, Immunity and Disease. *Pharmacological Reviews*, Pharmrev-Ar-2022–000618.

14. Chen, S., Yang, J., Wei, Y. & Wei, X. 2020. Epigenetic regulation of macrophages: from homeostasis maintenance to host defense. Cellular & Molecular Immunology, 17, 36–49.

15. Cheray, M. & Joseph, B. 2018. Epigenetics Control Microglia Plasticity. Front Cell Neurosci, 12, 243.

16. Cho, S. H., Chen, J. A., Sayed, F., Ward, M. E., Gao, F., Nguyen, T. A., Krabbe, G., Sohn, P. D., Lo, I., Minami, S., Devidze, N., Zhou, Y., Coppola, G. & Gan, L. 2015. SIRT1 deficiency in microglia contributes to cognitive decline in aging and neurodegeneration via epigenetic regulation of Il-1beta. J Neurosci, 35, 807–18.

17. Corley, M. J., Dye, C., D’antoni, M. L., Byron, M. M., Yo, K. L., Lum-Jones, A., Nakamoto, B., Valcour, V., Sahbandar, I., Shikuma, C. M., Ndhlovu, L. C. & Maunakea, A. K. 2016. Comparative Dna Methylation Profiling Reveals an Immunoepigenetic Signature of Hiv-related Cognitive Impairment. Sci Rep, 6, 33310.

18. Cull, A. H., Snetsinger, B., Buckstein, R., Wells, R. A. & Rauh, M. J. 2017. Tet2 restrains inflammatory gene expression in macrophages. Exp Hematol, 55, 56–70.e13.

19. Day, J. J., Childs, D., Guzman-Karlsson, M. C., Kibe, M., Moulden, J., Song, E., Tahir, A. & Sweatt, J. D. 2013. Dna methylation regulates associative reward learning. Nature Neuroscience, 16, 1445–1452.

20. De Carvalho, L. M., Wiers, C. E., Sun, H., Wang, G. J. & Volkow, N. D. 2021. Increased transcription of Tspo, HDAC2, and HDAC6 in the amygdala of males with alcohol use disorder. Brain Behav, 11, e01961.

21. Desplats, P., Dumaop, W., Cronin, P., Gianella, S., Woods, S., Letendre, S., Smith, D., Masliah, E. & Grant, I. 2014. Epigenetic alterations in the brain associated with Hiv-1 infection and methamphetamine dependence. PLos One, 9, e102555.

22. Doke, M., Pendyala, G. & Samikkannu, T. 2021. Psychostimulants and opioids differentially influence the epigenetic modification of histone acetyltransferase and histone deacetylase in astrocytes. Plos One, 16, e0252895.

23. Felger, J. C. & Treadway, M. T. 2017. Inflammation Effects on Motivation and Motor Activity: Role of Dopamine. Neuropsychopharmacology : official publication of the American College of Neuropsychopharmacology, 42, 216–241.

24. Feng, Y. & Lu, Y. 2021. Immunomodulatory Effects of Dopamine in Inflammatory Diseases. Front Immunol, 12, 663102.

25. Fukada, M., Nakayama, A., Mamiya, T., Yao, T.-P. & Kawaguchi, Y. 2016. Dopaminergic abnormalities in Hdac6-deficient mice. Neuropharmacology, 110, 470–479.

26. Galloway, A., Adeluyi, A., O’donovan, B., Fisher, M. L., Rao, C. N., Critchfield, P., Sajish, M., Turner, J. R. & Ortinski, P. I. 2018. Dopamine Triggers Ctcf-Dependent Morphological and Genomic Remodeling of Astrocytes. J Neurosci, 38, 4846–4858.

27. Gangisetty, O., Wynne, O., Jabbar, S., Nasello, C. & Sarkar, D. K. 2015. Fetal Alcohol Exposure Reduces Dopamine Receptor D2 and Increases Pituitary Weight and Prolactin Production via Epigenetic Mechanisms. PLos One, 10, e0140699.

28. Gao, Z., Lv, H., Wang, Y., Xie, Y., Guan, M. & Xu, Y. 2023. TET2 deficiency promotes anxiety and depression-like behaviors by activating NLRP3/Il-1β pathway in microglia of allergic rhinitis mice. Mol Med, 29, 160.

29. Gaskill, P. J., Carvallo, L., Eugenin, E. A. & Berman, J. W. 2012. Characterization and function of the human macrophage dopaminergic system: implications for Cns disease and drug abuse. J Neuroinflammation, 9, 203.

30. Gaskill, P. J., Yano, H. H., Kalpana, G. V., Javitch, J. A. & Berman, J. W. 2014. Dopamine receptor activation increases Hiv entry into primary human macrophages. PLos One, 9, e108232.

31. Ginno, P. A., Gaidatzis, D., Feldmann, A., Hoerner, L., Imanci, D., Burger, L., Zilbermann, F., Peters, A. H. F. M., Edenhofer, F., Smallwood, S. A., Krebs, A. R. & Schübeler, D. 2020. A genome-scale map of Dna methylation turnover identifies site-specific dependencies of Dnmt and Tet activity. Nature Communications, 11, 2680.

32. González, B., Gancedo, S. N., Garazatua, S. A. J., Roldán, E., Vitullo, A. D. & González, C. R. 2020. Dopamine Receptor D1 Contributes to Cocaine Epigenetic Reprogramming of Histone Modifications in Male Germ Cells. Front Cell Dev Biol, 8, 216.

33. González, B., Torres, O. V., Jayanthi, S., Gomez, N., Sosa, M. H., Bernardi, A., Urbano, F. J., García-Rill, E., Cadet, J. L. & Bisagno, V. 2019. The effects of single-dose injections of modafinil and methamphetamine on epigenetic and functional markers in the mouse medial prefrontal cortex: potential role of dopamine receptors. Prog Neuropsychopharmacol Biol Psychiatry, 88, 222–234.

34. Gopinath, A., Mackie, P., Hashimi, B., Buchanan, A. M., Smith, A. R., Bouchard, R., Shaw, G., Badov, M., Saadatpour, L., Gittis, A., Ramirez-Zamora, A., Okun, M. S., Streit, W. J., Hashemi, P. & Khoshbouei, H. 2022. Dat and Th expression marks human Parkinson’s disease in peripheral immune cells. Npj Parkinsons Dis, 8, 72.

35. Green, A. L., Eid, A., Zhan, L., Zarbl, H., Guo, G. L. & Richardson, J. R. 2019. Epigenetic Regulation of the Ontogenic Expression of the Dopamine Transporter. Front Genet, 10, 1099.

36. Hashimoto, K., Oreffo, R. O., Gibson, M. B., Goldring, M. B. & Roach, H. I. 2009. Dna demethylation at specific Cpg sites in the IL1b promoter in response to inflammatory cytokines in human articular chondrocytes. Arthritis Rheum, 60, 3303–13.

37. Hashimoto, K., Otero, M., Imagawa, K., De Andrés, M. C., Coico, J. M., Roach, H. I., Oreffo, R. O. C., Marcu, K. B. & Goldring, M. B. 2013. Regulated Transcription of Human Matrix Metalloproteinase 13 (MMP13) and Interleukin-1&#x3b2; (IL1b) Genes in Chondrocytes Depends on Methylation of Specific Proximal Promoter Cpg Sites *. Journal of Biological Chemistry, 288, 10061–10072.

38. He, X. B., Guo, F., Zhang, W., Fan, J., Le, W., Chen, Q., Ma, Y., Zheng, Y., Lee, S. H., Wang, H. J., Wu, Y., Zhou, Q. & Yang, R. 2024. JMJD3 deficiency disturbs dopamine biosynthesis in midbrain and aggravates chronic inflammatory pain. Acta Neuropathol Commun, 12, 201.

39. He, X. B., Yi, S. H., Rhee, Y. H., Kim, H., Han, Y. M., Lee, S. H., Lee, H., Park, C. H., Lee, Y. S., Richardson, E., Kim, B. W. & Lee, S. H. 2011. Prolonged membrane depolarization enhances midbrain dopamine neuron differentiation via epigenetic histone modifications. Stem Cells, 29, 1861–73.

40. Hillemacher, T., Rhein, M., Burkert, A., Heberlein, A., Wilhelm, J., Glahn, A., Muschler, M. A. N., Kahl, K. G., Kornhuber, J., Bleich, S. & Frieling, H. 2019. Dna-methylation of the dopamin receptor 2 gene is altered during alcohol withdrawal. Eur Neuropsychopharmacol, 29, 1250–1257.

41. Iwamoto, M., Nakamura, Y., Takemura, M., Hisaoka-Nakashima, K. & Morioka, N. 2020. TLR4-TAK1-p38 Mapk pathway and HDAC6 regulate the expression of sigma-1 receptors in rat primary cultured microglia. Journal of Pharmacological Sciences, 144, 23–29.

42. Jain, N., Shahal, T., Gabrieli, T., Gilat, N., Torchinsky, D., Michaeli, Y., Vogel, V. & Ebenstein, Y. 2019. Global modulation in Dna epigenetics during pro-inflammatory macrophage activation. Epigenetics, 14, 1183–1193.

43. Johnstone, A. L., Andrade, N. S., Barbier, E., Khomtchouk, B. B., Rienas, C. A., Lowe, K., Van Booven, D. J., Domi, E., Esanov, R., Vilca, S., Tapocik, J. D., Rodriguez, K., Maryanski, D., Keogh, M. C., Meinhardt, M. W., Sommer, W. H., Heilig, M., Zeier, Z. & Wahlestedt, C. 2021. Dysregulation of the histone demethylase KDM6b in alcohol dependence is associated with epigenetic regulation of inflammatory signaling pathways. Addict Biol, 26, e12816.

44. Kaminski, J. A., Schlagenhauf, F., Rapp, M., Awasthi, S., Ruggeri, B., Deserno, L., Banaschewski, T., Bokde, A. L. W., Bromberg, U., Büchel, C., Quinlan, E. B., Desrivières, S., Flor, H., Frouin, V., Garavan, H., Gowland, P., Ittermann, B., Martinot, J. L., Martinot, M. P., Nees, F., Orfanos, D. P., Paus, T., Poustka, L., Smolka, M. N., Fröhner, J. H., Walter, H., Whelan, R., Ripke, S., Schumann, G. & Heinz, A. 2018. Epigenetic variance in dopamine D2 receptor: a marker of Iq malleability? Transl Psychiatry, 8, 169.

45. Kashihara, K., Akiyama, K., Ishihara, T., Manabe, Y. & Abe, K. 1999. Differential regulation of Ap-1 Dna-binding activity by D1 and D2 dopamine receptor antagonists in the rat caudate-putamen and globus pallidus following a unilateral 6-Ohda lesion of the medial forebrain bundle. Neurol Res, 21, 175–9.

46. Kirchner, H., Nylen, C., Laber, S., Barrès, R., Yan, J., Krook, A., Zierath, J. R. & Näslund, E. 2014. Altered promoter methylation of PDK4, IL1 B, IL6, and Tnf after Roux-en Y gastric bypass. Surg Obes Relat Dis, 10, 671–8.

47. Kofman, S., Mohan, N., Sun, X., Ibric, L., Piermarini, E. & Qiang, L. 2022. Human mini brains and spinal cords in a dish: Modeling strategies, current challenges, and prospective advances. J Tissue Eng, 13, 20417314221113391.

48. Kohler, C. A., Freitas, T. H., Stubbs, B., Maes, M., Solmi, M., Veronese, N., De Andrade, N. Q., Morris, G., Fernandes, B. S., Brunoni, A. R., Herrmann, N., Raison, C. L., Miller, B. J., Lanctot, K. L. & Carvalho, A. F. 2018. Peripheral Alterations in Cytokine and Chemokine Levels After Antidepressant Drug Treatment for Major Depressive Disorder: Systematic Review and Meta-Analysis. Mol Neurobiol, 55, 4195–4206.

49. Kruidenier, L., Chung, C.-W., Cheng, Z., Liddle, J., Che, K., Joberty, G., Bantscheff, M., Bountra, C., Bridges, A., Diallo, H., Eberhard, D., Hutchinson, S., Jones, E., Katso, R., Leveridge, M., Mander, P. K., Mosley, J., Ramirez-Molina, C., Rowland, P., Schofield, C. J., Sheppard, R. J., Smith, J. E., Swales, C., Tanner, R., Thomas, P., Tumber, A., Drewes, G., Oppermann, U., Patel, D. J., Lee, K. & Wilson, D. M. 2012. A selective jumonji H3K27 demethylase inhibitor modulates the proinflammatory macrophage response. Nature, 488, 404–408.

50. Lepack, A. E., Werner, C. T., Stewart, A. F., Fulton, S. L., Zhong, P., Farrelly, L. A., Smith, A. C. W., Ramakrishnan, A., Lyu, Y., Bastle, R. M., Martin, J. A., Mitra, S., O’connor, R. M., Wang, Z.-J., Molina, H., Turecki, G., Shen, L., Yan, Z., Calipari, E. S., Dietz, D. M., Kenny, P. J. & Maze, I. 2020. Dopaminylation of histone H3 in ventral tegmental area regulates cocaine seeking. Science, 368, 197.

51. Lewis, C. R., Henderson-Smith, A., Breitenstein, R. S., Sowards, H. A., Piras, I. S., Huentelman, M. J., Doane, L. D. & Lemery-Chalfant, K. 2019. Dopaminergic gene methylation is associated with cognitive performance in a childhood monozygotic twin study. Epigenetics, 14, 310–323.

52. Li, H. D., Chen, X., Xu, J. J., Du, X. S., Yang, Y., Li, J. J., Yang, X. J., Huang, H. M., Li, X. F., Wu, M. F., Zhang, C., Zhang, C., Li, Z., Wang, H., Meng, X. M., Huang, C. & Li, J. 2020. DNMT3b-mediated methylation of ZSWIM3 enhances inflammation in alcohol-induced liver injury via regulating TRAF2-mediated Nf-κb pathway. Clin Sci (Lond), 134, 1935–1956.

53. Li, L. C. & Dahiya, R. 2002. MethPrimer: designing primers for methylation PCRs. Bioinformatics, 18, 1427–31.

54. Li, Y., Rong, J., Zhong, H., Liang, M., Zhu, C., Chang, F. & Zhou, R. 2021. Prenatal Stress Leads to the Altered Maturation of Corticostriatal Synaptic Plasticity and Related Behavioral Impairments Through Epigenetic Modifications of Dopamine D2 Receptor in Mice. Mol Neurobiol, 58, 317–328.

55. Liu, L., Wu, Y., Wang, B., Jiang, Y., Lin, L., Li, X. & Yang, S. 2021. Da-DRD5 signaling controls colitis by regulating colonic M1/M2 macrophage polarization. Cell Death & Disease, 12, 500.

56. Liu, S., Chen, X., Chen, R., Wang, J., Zhu, G., Jiang, J., Wang, H., Duan, S. & Huang, J. 2017. Diagnostic role of Wnt pathway gene promoter methylation in non small cell lung cancer. Oncotarget, 8, 36354–36367.

57. Mackie, P. M., Gopinath, A., Montas, D. M., Nielsen, A., Smith, A., Nolan, R. A., Runner, K., Matt, S. M., Mcnamee, J., Riklan, J. E., Adachi, K., Doty, A., Ramirez-Zamora, A., Yan, L., Gaskill, P. J., Streit, W. J., Okun, M. S. & Khoshbouei, H. 2022. Functional characterization of the biogenic amine transporters on human macrophages. Jci Insight.

58. Marei, H. E. 2025. Epigenetic Editing in Neurological and Neuropsychiatric Disorders: Pioneering Next-Gen Therapeutics for Precision Gene Control. Mol Neurobiol, 63, 330.

59. Marion-Poll, L., Roussarie, J. P., Taing, L., Dard-Dascot, C., Servant, N., Jaszczyszyn, Y., Jordi, E., Mulugeta, E., Hervé, D., Bourc’his, D., Greengard, P., Thermes, C. & Girault, J. A. 2022. Dna methylation and hydroxymethylation characterize the identity of D1 and D2 striatal projection neurons. Commun Biol, 5, 1321.

60. Matt, S. M. & Gaskill, P. J. 2019. Where Is Dopamine and how do Immune Cells See it?: Dopamine-Mediated Immune Cell Function in Health and Disease. J Neuroimmune Pharmacol.

61. Matt, S. M., Lawson, M. A. & Johnson, R. W. 2016. Aging and peripheral lipopolysaccharide can modulate epigenetic regulators and decrease Il-1beta promoter Dna methylation in microglia. Neurobiol Aging, 47, 1–9.

62. Matt, S. M., Nickoloff-Bybel, E. A., Rong, Y., Runner, K., Johnson, H., O’connor, M. H., Haddad, E. K. & Gaskill, P. J. 2021. Dopamine Levels Induced by Substance Abuse Alter Efficacy of Maraviroc and Expression of CCR5 Conformations on Myeloid Cells: Implications for Neurohiv. Front Immunol, 12, 663061.

63. Matt, S. M., Nolan, R., Manikandan, S., Agarwal, Y., Channer, B., Oteju, O., Daniali, M., Canagarajah, J. A., Lupone, T., Mompho, K., Runner, K., Nickoloff-Bybel, E., Li, B., Niu, M., Schlachetzki, J. C. M., Fox, H. S. & Gaskill, P. J. 2025. Dopamine-driven increase in Il-1β in myeloid cells is mediated by differential dopamine receptor expression and exacerbated by Hiv. Journal of Neuroinflammation, 22, 91.

64. Melka, M. G., Castellani, C. A., Laufer, B. I., Rajakumar, R. N., O’reilly, R. & Singh, S. M. 2013. Olanzapine induced Dna methylation changes support the dopamine hypothesis of psychosis. J Mol Psychiatry, 1, 19.

65. Muench, C., Wiers, C. E., Cortes, C. R., Momenan, R. & Lohoff, F. W. 2018. Dopamine Transporter Gene Methylation is Associated with Nucleus Accumbens Activation During Reward Processing in Healthy but not Alcohol-Dependent Individuals. Alcohol Clin Exp Res, 42, 21–31.

66. Nestler, E. J. & Lüscher, C. 2019. The Molecular Basis of Drug Addiction: Linking Epigenetic to Synaptic and Circuit Mechanisms. Neuron, 102, 48–59.

67. Netea, M. G., Joosten, L. A. B., Latz, E., Mills, K. H. G., Natoli, G., Stunnenberg, H. G., O’neill, L. A. J. & Xavier, R. J. 2016. Trained immunity: A program of innate immune memory in health and disease. Science, 352.

68. Nickoloff-Bybel, E. A., Mackie, P., Runner, K., Matt, S. M., Khoshbouei, H. & Gaskill, P. J. 2019. Dopamine increases Hiv entry into macrophages by increasing calcium release via an alternative signaling pathway. Brain Behav Immun, 82, 239–252.

69. Nolan, R. A., Muir, R., Runner, K., Haddad, E. K. & Gaskill, P. J. 2018. Role of Macrophage Dopamine Receptors in Mediating Cytokine Production: Implications for Neuroinflammation in the Context of Hiv-Associated Neurocognitive Disorders. Journal of Neuroimmune Pharmacology.

70. Nolan, R. A., Reeb, K. L., Rong, Y., Matt, S. M., Johnson, H. S., Runner, K. & Gaskill, P. J. 2020. Dopamine activates Nf-κb and primes the NLRP3 inflammasome in primary human macrophages. *Brain, Behavior*, & Immunity - Health, 2, 100030.

71. Ookubo, M., Kanai, H., Aoki, H. & Yamada, N. 2013. Antidepressants and mood stabilizers effects on histone deacetylase expression in C57bl/6 mice: Brain region specific changes. J Psychiatr Res, 47, 1204–14.

72. Penrod, R. D., Carreira, M. B., Taniguchi, M., Kumar, J., Maddox, S. A. & Cowan, C. W. 2018. Novel role and regulation of HDAC4 in cocaine-related behaviors. Addict Biol, 23, 653–664.

73. Rossetti, M. F., Schumacher, R., Gastiazoro, M. P., Lazzarino, G. P., Andreoli, M. F., Stoker, C., Varayoud, J. & Ramos, J. G. 2020. Epigenetic Dysregulation of Dopaminergic System by Maternal Cafeteria Diet During Early Postnatal Development. Neuroscience, 424, 12–23.

74. Rubino, A., D’addario, C., Di Bartolomeo, M., Michele Salamone, E., Locuratolo, N., Fattapposta, F., Vanacore, N. & Pascale, E. 2020. Dna methylation of the 5’-Utr Dat 1 gene in Parkinson’s disease patients. Acta Neurol Scand, 142, 275–280.

75. Sasagawa, T., Horii-Hayashi, N., Okuda, A., Hashimoto, T., Azuma, C. & Nishi, M. 2017. Long-term effects of maternal separation coupled with social isolation on reward seeking and changes in dopamine D1 receptor expression in the nucleus accumbens via Dna methylation in mice. Neurosci Lett, 641, 33–39.

76. Schroeder, F. A., Penta, K. L., Matevossian, A., Jones, S. R., Konradi, C., Tapper, A. R. & Akbarian, S. 2008. Drug-induced activation of dopamine D(1) receptor signaling and inhibition of class I/Ii histone deacetylase induce chromatin remodeling in reward circuitry and modulate cocaine-related behaviors. Neuropsychopharmacology, 33, 2981–92.

77. Sharma, D., Malik, A., Locher, V., Mcgrath, S., Zabala, S., Grover, H., Ciszewski, C., Chen, L., Yang, D., Chiu, I. M. & Jabri, B. 2025. A myeloid Tet2-Il-1β axis modulates intestinal inflammation by restricting catecholaminergic stimulation of enterochromaffin cell differentiation. Immunity, 58, 2785–2798.e4.

78. Shi, F. D. & Yong, V. W. 2025. Neuroinflammation across neurological diseases. Science, 388, eadx0043.

79. Södersten, E., Feyder, M., Lerdrup, M., Gomes, A. L., Kryh, H., Spigolon, G., Caboche, J., Fisone, G. & Hansen, K. 2014. Dopamine signaling leads to loss of Polycomb repression and aberrant gene activation in experimental parkinsonism. PLos Genet, 10, e1004574.

80. Staes, N., White, C. M., Guevara, E. E., Eens, M., Hopkins, W. D., Schapiro, S. J., Stevens, J. M. G., Sherwood, C. C. & Bradley, B. J. 2022. Chimpanzee Extraversion scores vary with epigenetic modification of dopamine receptor gene D2 (DRD2) and early rearing conditions. Epigenetics, 17, 1701–1714.

81. Suchanecka, A., Rexław, R., Chmielowiec, K., Chmielowiec, J., Masiak, J. & Grzywacz, A. 2025. Methylation Status of the DAT1 Dopamine Transporter Gene in Individuals With Cannabis Use Disorder: Associations With Personality Traits. Genes Brain Behav, 24, e70040.

82. Sun, X., Kofman, S., Ogbolu, V. C., Karch, C. M., Ibric, L. & Qiang, L. 2024. Vascularized Brain Assembloids With Enhanced Cellular Complexity Provide Insights Into the Cellular Deficits of Tauopathy. Stem Cells, 42, 107–115.

83. Taniguchi, M., Carreira, M. B., Smith, L. N., Zirlin, B. C., Neve, R. L. & Cowan, C. W. 2012. Histone deacetylase 5 limits cocaine reward through camp-induced nuclear import. Neuron, 73, 108–20.

84. Tekpli, X., Landvik, N. E., Anmarkud, K. H., Skaug, V., Haugen, A. & Zienolddiny, S. 2013. Dna methylation at promoter regions of interleukin 1b, interleukin 6, and interleukin 8 in non-small cell lung cancer. Cancer Immunol Immunother, 62, 337–45.

85. Thomas Broome, S., Louangaphay, K., Keay, K. A., Leggio, G. M., Musumeci, G. & Castorina, A. 2020. Dopamine: an immune transmitter. Neural Regeneration Research, 15.

86. Torquet, N., Marti, F., Campart, C., Tolu, S., Nguyen, C., Oberto, V., Benallaoua, M., Naudé, J., Didienne, S., Debray, N., Jezequel, S., Le Gouestre, L., Hannesse, B., Mariani, J., Mourot, A. & Faure, P. 2018. Social interactions impact on the dopaminergic system and drive individuality. Nature Communications, 9, 3081.

87. Unland, R., Kerl, K., Schlosser, S., Farwick, N., Plagemann, T., Lechtape, B., Clifford, S. C., Kreth, J. H., Gerss, J., Mühlisch, J., Richter, G. H., Hasselblatt, M. & Frühwald, M. C. 2014. Epigenetic repression of the dopamine receptor D4 in pediatric tumors of the central nervous system. J Neurooncol, 116, 237–49.

88. Van Der Heijden, C., Groh, L., Keating, S. T., Kaffa, C., Noz, M. P., Kersten, S., Van Herwaarden, A. E., Hoischen, A., Joosten, L. A. B., Timmers, H., Netea, M. G. & Riksen, N. P. 2020. Catecholamines Induce Trained Immunity in Monocytes In Vitro and In Vivo. Circ Res, 127, 269–283.

89. Vento-Tormo, R., Álvarez-Errico, D., Garcia-Gomez, A., Hernández-Rodríguez, J., Buán, S., Basagaña, M., Méndez, M., Yagüe, J., Juan, M., ARóstegui, J. I. & Ballestar, E. 2017. Dna demethylation of inflammasome-associated genes is enhanced in patients with cryopyrin-associated periodic syndromes. J Allergy Clin Immunol, 139, 202–211.e6.

90. Vilca, S., Wahlestedt, C., Izenwasser, S., Gainetdinov, R. R. & Pardo, M. 2023. Dopamine Transporter Knockout Rats Display Epigenetic Alterations in Response to Cocaine Exposure. Biomolecules, 13.

91. Vucetic, Z., Carlin, J. L., Totoki, K. & Reyes, T. M. 2012. Epigenetic dysregulation of the dopamine system in diet-induced obesity. J Neurochem, 120, 891–8.

92. Wang, H., Divaris, K., Pan, B., Li, X., Lim, J.-H., Saha, G., Barovic, M., Giannakou, D., Korostoff, J. M., Bing, Y., Sen, S., Moss, K., Wu, D., Beck, J. D., Ballantyne, C. M., Natarajan, P., North, K. E., Netea, M. G., Chavakis, T. & Hajishengallis, G. 2024. Clonal hematopoiesis driven by mutated DNMT3a promotes inflammatory bone loss. Cell, 187, 3690–3711.e19.

93. Wang, J., Hodes, G. E., Zhang, H., Zhang, S., Zhao, W., Golden, S. A., Bi, W., Menard, C., Kana, V., Leboeuf, M., Xie, M., Bregman, D., Pfau, M. L., Flanigan, M. E., Esteban-FERNández, A., Yemul, S., Sharma, A., Ho, L., Dixon, R., Merad, M., Han, M. H., Russo, S. J. & Pasinetti, G. M. 2018. Epigenetic modulation of inflammation and synaptic plasticity promotes resilience against stress in mice. Nat Commun, 9, 477.

94. Weinmann, A. S., Mitchell, D. M., Sanjabi, S., Bradley, M. N., Hoffmann, A., Liou, H. C. & Smale, S. T. 2001. Nucleosome remodeling at the Il-12 p40 promoter is a Tlr-dependent, Rel-independent event. Nat Immunol, 2, 51–7.

95. Wiers, C. E., Lohoff, F. W., Lee, J., Muench, C., Freeman, C., Zehra, A., Marenco, S., Lipska, B. K., Auluck, P. K., Feng, N., Sun, H., Goldman, D., Swanson, J. M., Wang, G. J. & Volkow, N. D. 2018. Methylation of the dopamine transporter gene in blood is associated with striatal dopamine transporter availability in Adhd: A preliminary study. Eur J Neurosci, 48, 1884–1895.

96. Wu, C., Li, A., Hu, J. & Kang, J. 2019. Histone deacetylase 2 is essential for Lps-induced inflammatory responses in macrophages. Immunol Cell Biol, 97, 72–84.

97. Wu, T., Cai, W. & Chen, X. 2023. Epigenetic regulation of neurotransmitter signaling in neurological disorders. Neurobiol Dis, 184, 106232.

98. Wu, X., Chen, P. S., Dallas, S., Wilson, B., Block, M. L., Wang, C. C., Kinyamu, H., Lu, N., Gao, X., Leng, Y., Chuang, D. M., Zhang, W., Lu, R. B. & Hong, J. S. 2008. Histone deacetylase inhibitors up-regulate astrocyte Gdnf and Bdnf gene transcription and protect dopaminergic neurons. Int J Neuropsychopharmacol, 11, 1123–34.

99. Xiao, J., Li, Y., Prandovszky, E., Karuppagounder, S. S., Talbot, C. C., Jr., Dawson, V. L., Dawson, T. M. & Yolken, R. H. 2014. Microrna-132 dysregulation in Toxoplasma gondii infection has implications for dopamine signaling pathway. Neuroscience, 268, 128–38.

100. Yan, B., Xie, S., Liu, Y., Liu, W., Li, D., Liu, M., Luo, H. R. & Zhou, J. 2018. Histone deacetylase 6 modulates macrophage infiltration during inflammation. Theranostics, 8, 2927–2938.

101. Yan, S., Wei, X., Jian, W., Qin, Y., Liu, J., Zhu, S., Jiang, F., Lou, H. & Zhang, B. 2020. Pharmacological Inhibition of HDAC6 Attenuates NLRP3 Inflammatory Response and Protects Dopaminergic Neurons in Experimental Models of Parkinson’s Disease. Front Aging Neurosci, 12, 78.

102. Yan, Y., Jiang, W., Liu, L., Wang, X., Ding, C., Tian, Z. & Zhou, R. 2015. Dopamine Controls Systemic Inflammation through Inhibition of NLRP3 Inflammasome. Cell, 160, 62–73.

103. Yu, J., Qiu, Y., Yang, J., Bian, S., Chen, G., Deng, M., Kang, H. & Huang, L. 2016. DNMT1-PPARγ pathway in macrophages regulates chronic inflammation and atherosclerosis development in mice. Scientific Reports, 6, 30053.

104. Zhang, L., Lou, D., Jiao, H., Zhang, D., Wang, X., Xia, Y., Zhang, J. & Xu, M. 2004. Cocaine-induced intracellular signaling and gene expression are oppositely regulated by the dopamine D1 and D3 receptors. J Neurosci, 24, 3344–54.

105. Zhang, Q., Zhao, K., Shen, Q., Han, Y., Gu, Y., Li, X., Zhao, D., Liu, Y., Wang, C., Zhang, X., Su, X., Liu, J., Ge, W., Levine, R. L., Li, N. & Cao, X. 2015a. Tet2 is required to resolve inflammation by recruiting Hdac2 to specifically repress Il-6. Nature, 525, 389–393.

106. Zhang, W., Jiang, J., Xu, Z., Yan, H., Tang, B., Liu, C., Chen, C. & Meng, Q. 2023a. Microglia-containing human brain organoids for the study of brain development and pathology. Mol Psychiatry, 28, 96–107.

107. Zhang, Y., Gao, Y., Ding, Y., Jiang, Y., Chen, H., Zhan, Z. & Liu, X. 2023b. Targeting KAT2a inhibits inflammatory macrophage activation and rheumatoid arthritis through epigenetic and metabolic reprogramming. MedComm, 4, e306.

108. Zhang, Y., Wang, Y., Wang, L., Bai, M., Zhang, X. & Zhu, X. 2015b. Dopamine Receptor D2 and Associated microRNAs Are Involved in Stress Susceptibility and Resistance to Escitalopram Treatment. Int J Neuropsychopharmacol, 18.

109. Zhu, H., Wang, G. & Qian, J. 2016. Transcription factors as readers and effectors of Dna methylation. Nat Rev Genet, 17, 551–65.

